# Co-translational sorting enables a single mRNA to generate distinct polysomes with different localizations and protein fates

**DOI:** 10.1101/2024.03.20.585881

**Authors:** Soha Salloum, Martial Séveno, Khadija El Koulali, Stephanie Rialle, Simon George, Benedicte Lemmers, Kazem Zibara, Carolina Eliscovich, Michael Hahne, Edouard Bertrand

## Abstract

β-catenin is a multi-functional protein playing essential roles in tissue homeostasis and cancer. It bridges E-cadherin to the cytoskeleton and also activates transcription in response to Wnt. Plasma membrane β-catenin is stable whereas without Wnt, cytoplasmic β-catenin is degraded by the destruction complex, composed of APC and Axin. Here, we show that APC and Axin associate with many mRNAs and that this occurs via the nascent protein chains. Notably, APC and Axin bind β-catenin mRNAs present as either single polysome or polysome condensates, and co-translational interactions constitute the major fraction of their binding to the β-catenin protein. Remarkably, E-cadherin also binds β-catenin co-translationally, and β-catenin mRNAs localize either with APC in the cytosol or E-cadherin at the plasma membrane. Thus, co-translational interactions sort β-catenin mRNAs into distinct polysome populations that spatially segregate in cells and synthesize proteins with different functions. Co-translational polysome sorting provides a mechanism to regulate the fate of multi-functional proteins.

## Introduction

The Wnt/β-catenin signalling pathway is evolutionary conserved and its alterations leads to cancer (Gergen and Wieschaus, 1986; Mccrea et al., 1991, reviewed in Bian et al., 2020; Kwong et al., 2009; Nakanishi et al., 2020). The role of β-catenin in cancer was discovered through the study of familial adenomatous polyposis, a hereditary form of colorectal cancer (Kwong et al., 2009). It was found that APC is the main causative gene of the disease and that APC directly binds β-catenin and downregulates its cellular levels (Kinzler et al., 1991; Groden et al., 1991 Grainger and Willert, 2018; MacDonald et al., 2009; Stamos and Weis, 2013). Since then, the Wnt/β-catenin pathway has been shown to be important not only for colorectal tumors (CRC), but also for many other tumor types, including breast, lung, liver and hematopoietic malignancies (Nusse and Varmus, 1982, Miyoshi et al., 1998; Kwong et al., 2009). Aberrant activation of the Wnt pathway leads to the transcriptional activation of target genes by β-catenin (Bian et al., 2020; Grainger and Willert, 2018; MacDonald et al., 2009; Stamos and Weis, 2013). Over the years, many components of the Wnt/β-catenin pathway have been identified as being important in tumorigenesis, ranging from genes involved in Wnt secretion to its transcriptional response (Morin et al., 1997; Korinek et al., 1998; Bian et al., 2020; Nakanishi et al., 2020). The canonical Wnt pathway involves many Wnt ligands and receptors, but all converge to β-catenin, highlighting its central role in the pathway (Bian et al., 2020; Grossmann et al., 2012). Remarkably, β-catenin is hyperactivated in nearly all CRC tumors, and this appears to be an early and key event during tumorigenesis (Iwao et al., 1998; Polakis et al., 2007; Bian et al., 2020; Kwong et al., 2009). About 80% of CRC have mutations in APC, and another 5-10% have mutations in other components of the Wnt/β-catenin pathway (Miyoshi et al., 1992; Bian et al., 2020; Schaefer and Peifer, 2019). Deciphering the regulation of β-catenin and the function of APC is thus essential for understanding the biology of CRC, and major efforts have been made in this direction.

The β-catenin protein is mainly regulated at the level of its stability. In presence of Wnt, β-catenin is stabilized and translocated to the nucleus to activate transcription of target genes (Behrens et al., 1996; Bian et al., 2020; Grainger and Willert, 2018; Grossmann et al., 2012; MacDonald et al., 2009; Stamos and Weis, 2013; Schaefer and Peifer, 2019). In absence of Wnt, however, β-catenin is rapidly degraded by the proteasome (Aberle et al., 1997). The main factors involved in β-catenin degradation are Axin, APC, and the kinases CK1α and GSK3, which altogether form the “destruction complex” (Rubinfeld et al., 1996; Ikeda et al., 1998; Bian et al., 2020; Grainger and Willert, 2018; MacDonald et al., 2009; Stamos and Weis, 2013). In absence of Wnt, the destruction complex binds β-catenin and targets it for degradation. In presence of Wnt, this complex becomes recruited to the Wnt receptor at the plasma membrane via the Dishevelled protein that heteromerizes with Axin. There, GSK3 is inhibited by the co-receptors LRP5/6 and the destruction complex can no longer access cytoplasmic β-catenin (Cselenyi et al., 2008; Piao et al., 2008, Bian et al., 2020; Grainger and Willert, 2018; Grossmann et al., 2012; MacDonald et al., 2009; Stamos and Weis, 2013; Schaefer and Peifer, 2019).

Axin is an essential player of the destruction complex and it acts as a scaffolding protein (Hart et al., 1998; Bian et al., 2020; Grainger and Willert, 2018; MacDonald et al., 2009; Stamos and Weis, 2013). It binds β-catenin as well as APC, CK1α and GSK3. It promotes β-catenin phosphorylation by bringing β-catenin in proximity to the kinases CK1α and GSK3 (Liu et al., 1999, Liu et al., 2002; Bian et al., 2020; Grainger and Willert, 2018; MacDonald et al., 2009; Stamos and Weis, 2013).Phosphorylation by GSK3 on S33, S37, T41 and S45 leads to β-catenin recognition by the E3 ubiquitin ligase bTrCP, which targets it for proteasomal degradation (Aberle et al., 1997; Hart et al., 1998). APC is a very large protein that also functions as a scaffold. It occurs as a dimer and each monomer has tens of binding sites for β-catenin (Grainger and Willert, 2018; Grossmann et al., 2012; MacDonald et al., 2009; Stamos and Weis, 2013; Schaefer and Peifer, 2019). Interestingly, Axin can make oligomers, and APC can multimerize with these oligomers to make large molecular (Schaefer and Peifer, 2019). Despite a very large number of studies and an intense scrutiny, it is important to note that the exact role of APC in β-catenin degradation is still poorly understood. In particular, APC can be dispensable to trigger β-catenin phosphorylation and proteasomal targeting, for instance when Axin is present in high concentration (Grainger and Willert, 2018; MacDonald et al., 2009; Stamos and Weis, 2013). Given the stoichiometry of the APC/β-catenin interaction (1 APC binds tens of β-catenin), it is believed that APC works as a β-catenin “sink”, capturing free molecules and handling them to Axin for degradation (Grossmann et al., 2012; MacDonald et al., 2009; Stamos and Weis, 2013; Schaefer and Peifer, 2019).

Many proteins are present in multiple molecular complexes that can have distinct, unrelated functions. β-catenin is a very good example of multifunctional proteins, since it is present in several molecular complexes that have different localizations and functions (Peifer, 1993; Bian et al., 2020; Grainger and Willert, 2018; Grossmann et al., 2012; MacDonald et al., 2009; Stamos and Weis, 2013). First, β-catenin is the main transcription factor of the Wnt pathway. In this case, it localizes in the nucleus and associates with LEF/TCF transcription factors (Graham et al., 2000; Bian et al., 2020; Kwong et al., 2009. Second, β-catenin binds APC and Axin in the cytoplasm, and this targets it to proteasomal degradation. Finally, β-catenin links adherens junctions with the actin cytoskeleton. Here, it localizes at the plasma membrane and it makes a complex with α-catenin, F-actin and E-cadherin (Drees et al., 2005; Huber and Weis, 2001; Ozawa et al., 1989). The interactions of β-catenin with APC/Axin, E-cadherin and LEF/TCF are all mutually exclusive, providing a simple mechanism to separate β-catenin functions.

In this study, we explored the links between β-catenin mRNAs and the destruction complex. We show that APC and Axin bind a variety of mRNAs including the one coding for β-catenin, and we observe that this occurs via the nascent protein chains. Remarkably, β-catenin makes co-translational interactions not only with the destruction complex but also with E-cadherin, and these mutually exclusive interactions sort β-catenin mRNAs into two populations of polysomes that localize differently within the cell and synthesize proteins with different fates and functions. We refer to this mechanism as polysome sorting. It provides an elegant way to regulate separately in space and time the functions of multifunctional proteins.

## Results

### APC binds mRNAs in a translation-dependent manner

We recently performed a screen to characterize intra-cellular mRNA localization and local translation (Chouaib et al., 2020). In this screen, we used a collection of HeLa cell lines, each containing a bacterial artificial chromosome (BAC) carrying an entire human gene with a GFP tag inserted at the N- or C-terminus of the corresponding protein. In the case of the β-catenin-GFP BAC cell line, we observed that the β-catenin mRNA formed cytoplasmic condensates containing multiple mRNA molecules as well as APC and Axin (Chouaib et al., 2020; see Figure 1A). The mRNA foci were thus proposed to be specialized translation factories whose function is to target β-catenin for degradation.

**Figure 1:**
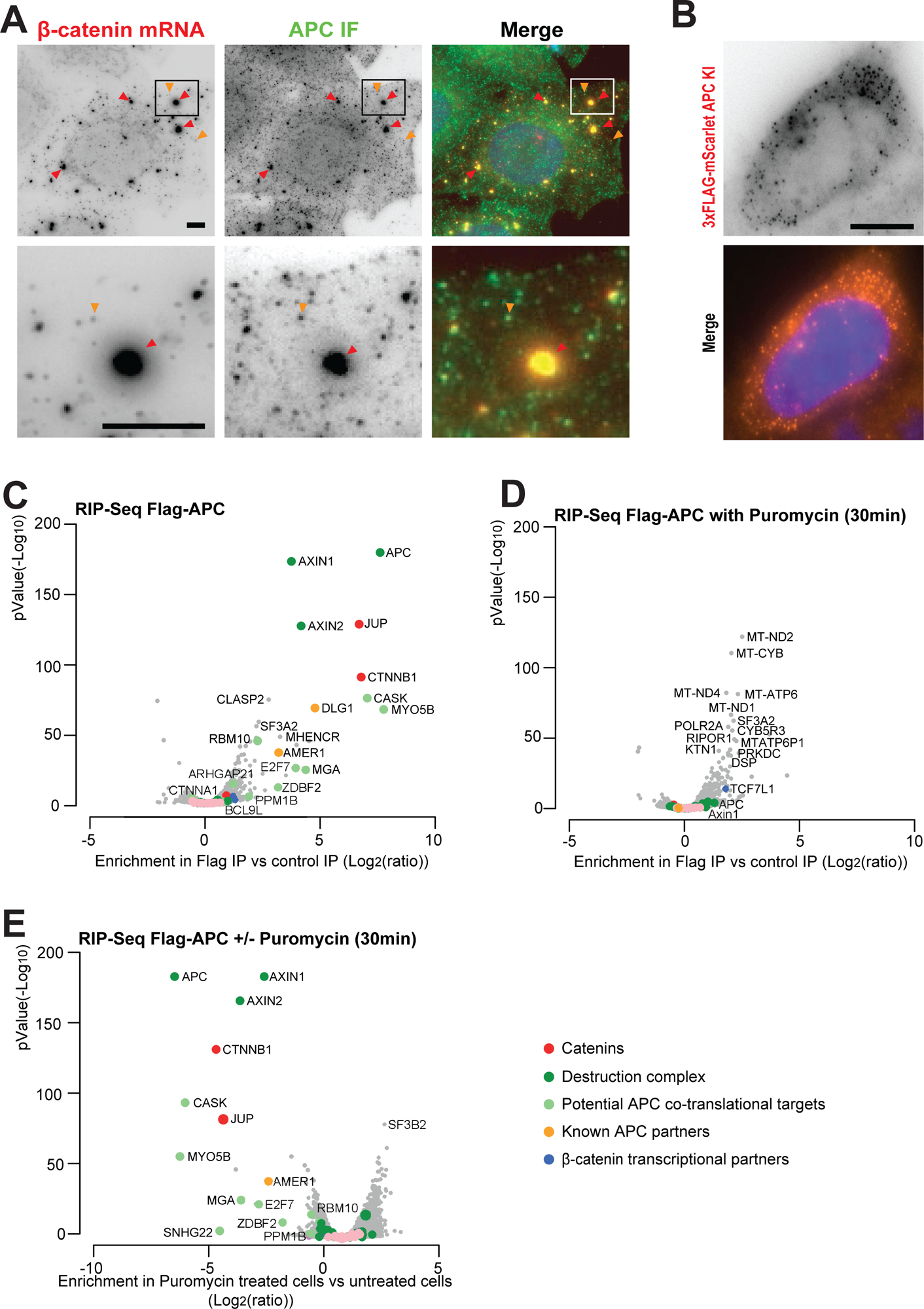
APC interacts with RNAs in a translation-dependent manner. **A-**Images are micrographs of HeLa β-catenin GFP BAC cells. Left and red: Cy3 fluorescent signals corresponding to β-catenin mRNAs detected with smiFISH; middle and green: FITC signals corresponding to APC detected by immunofluorescence. For each cell, a zoom of the boxed area is shown below. Red arrows show APC and β-catenin mRNA foci; orange arrows show single β-catenin mRNAs. DNA is stained with DAPI. Scale bar: 5 microns. **B**-Images are micrographs of HEK293 3xFLAG-mScarlet-APC knock-in cells. Left and red: 3xFLAG-mScarlet-APC signals. DNA is stained with DAPI. Scale bar: 10 microns. **C**-Volcano plots depicting the interaction of 3xFLAG-mScarlet-APC with RNAs in HEK293 knock-in cells. The x axis represents the enrichment (log_2_ [ratio]) of the RNAs immunoprecipitated with anti-FLAG antibody conjugated beads (FLAG-IP) compared to beads without FLAG antibody (control IP). The y axis represents significance, displayed as -log10 (p-Value). Experiments were done in duplicates and statistical significance is considered for p-Value <0.01. The colored points in the plots represent different categories of proteins encoded by the identified mRNAs. The categories are listed at the bottom right. **D**-As in C except that cells were treated with puromycin for 30 min at 100 μg/ml. **E**-Volcano plot comparing RNAs immunoprecipitated in the FLAG IPs, in puromycin *versus* untreated condition. Legend as in C.

To determine more directly whether APC associates with β-catenin mRNAs, we used HEK293 cells as a model system. These cells are easy to manipulate and they have often been used to study the Wnt pathway as it is functional in this cell line (Jia et al., 2008). We introduced a 3xFLAG-mScarlet tag at the N-terminal region of APC by CRISPR genome editing (Figure S1A), thereby preserving the endogenous APC expression levels. Heterozygous clones carrying a tagged APC allele were verified by PCR genotyping and clone N112 was kept for further study (Figure S1B). Fluorescent microscopy revealed that 3xFLAG-mScarlet-APC was diffusely distributed in the cytoplasm and formed puncta in some cells, as expected from previous studies (Figure 1B; Chouaib et al., 2020). We performed a RIP-seq experiment and immunoprecipitated (IP’ed) APC using anti-FLAG beads and beads without antibody as a control, and sequenced the associated RNAs. APC interacted strongly and significantly with β-catenin mRNA (CTNNB1), as well as a number of other mRNAs (Figure 1C, Table S1). Some of these mRNAs encoded direct protein partners of APC such as Axin1, Axin2, DLG1 (Matsumine et al., 1996), AMER1 (Grohmann et al., 2007) and JUP, which encodes the ϕ3-catenin paralog γ-catenin (Rubinfeld et al., 1995), while others encoded proteins identified as potential APC partners in large-scale screens such as CASK (Buljan et al., 2020) and ARHGAP21 (Hein et al., 2015). In addition, APC interacted with mRNAs encoding proteins that were not previously described as APC partners, such as MYO5B, E2F7, RBM10, ZDBF2 and MGA (Figure 1C).

APC has been previously shown to directly interact with mRNAs by CLIP-seq (Preitner et al., 2014), and in particular β-catenin. It also directly binds polyA+ mRNAs in crosslinking-based screens for RNA binding proteins (Trendel et al., 2019). One possibility that explains these results would be that APC is a non-canonical RNA binding protein. Another possibility would be that APC binds these mRNA by interacting with the nascent protein chain emerging from the ribosome, whose proximity to mRNAs would then enable its cross-linking. To discriminate between these possibilities, we repeated the RIP-seq experiments in cells treated with puromycin, which induces translation termination and detaches ribosomes from mRNAs. Remarkably, the mRNA associated with APC in untreated cells were no longer interacting after 30 minutes of puromycin treatment, including β-catenin (CTNNB1), Axin1, Axin2, CASK, JUP, MYO5B and AMER1 mRNAs (Figure 1D-E; S1C; Table S1). Some transcripts remained bound to APC in puromycin-treated cells, and in particular mitochondrial mRNAs (Figure 1D), possibly reflecting an association of APC with this organelle (Mills et al., 2016). Our results thus confirmed that APC associates with specific mRNAs including β- and γ-catenin, and demonstrated that these interactions occur co-translationally via the nascent protein chain.

### APC partners can be strictly co-translational, co- and post-translational, or strictly post-translational

Because the association of APC with many mRNAs was translation dependent, we next aimed to compare the protein and mRNA partners of APC. We thus examined APC protein interactions using quantitative label-free proteomics. We performed the same FLAG and control IPs as above, using our 3xFLAG-mScarlet-APC HEK293 cell line, and analyzed the pelleted proteins by mass spectrometry (Figure 2A, Table S2). As expected, we identified known APC protein partners specifically enriched in the FLAG IP, including as β-catenin (CTNNB1), α-catenin (CTNNA1), AMER1, CASK, DLG1 and CTBP1/CTBP2, two related proteins involved in transcription repression (Sollerbrant et al., 1996). Surprisingly, two members of the CTLH complex were significantly enriched in the FLAG IP, RANBP9 and WDR26, while other members of this complex were also enriched but with lower significance (i.e. GID8, MAEA YPEL5, ARMC8, MKLN1). The CTLH complex is an ubiquitin ligase analogous to the GID complex in yeast, and it was not known to associate with APC although interactions between individual members of this complex and components of the Wnt/β-catenin pathway were previously reported (Goto et al., 2016; Liang et al., 2017; Sato et al., 2020).

**Figure 2:**
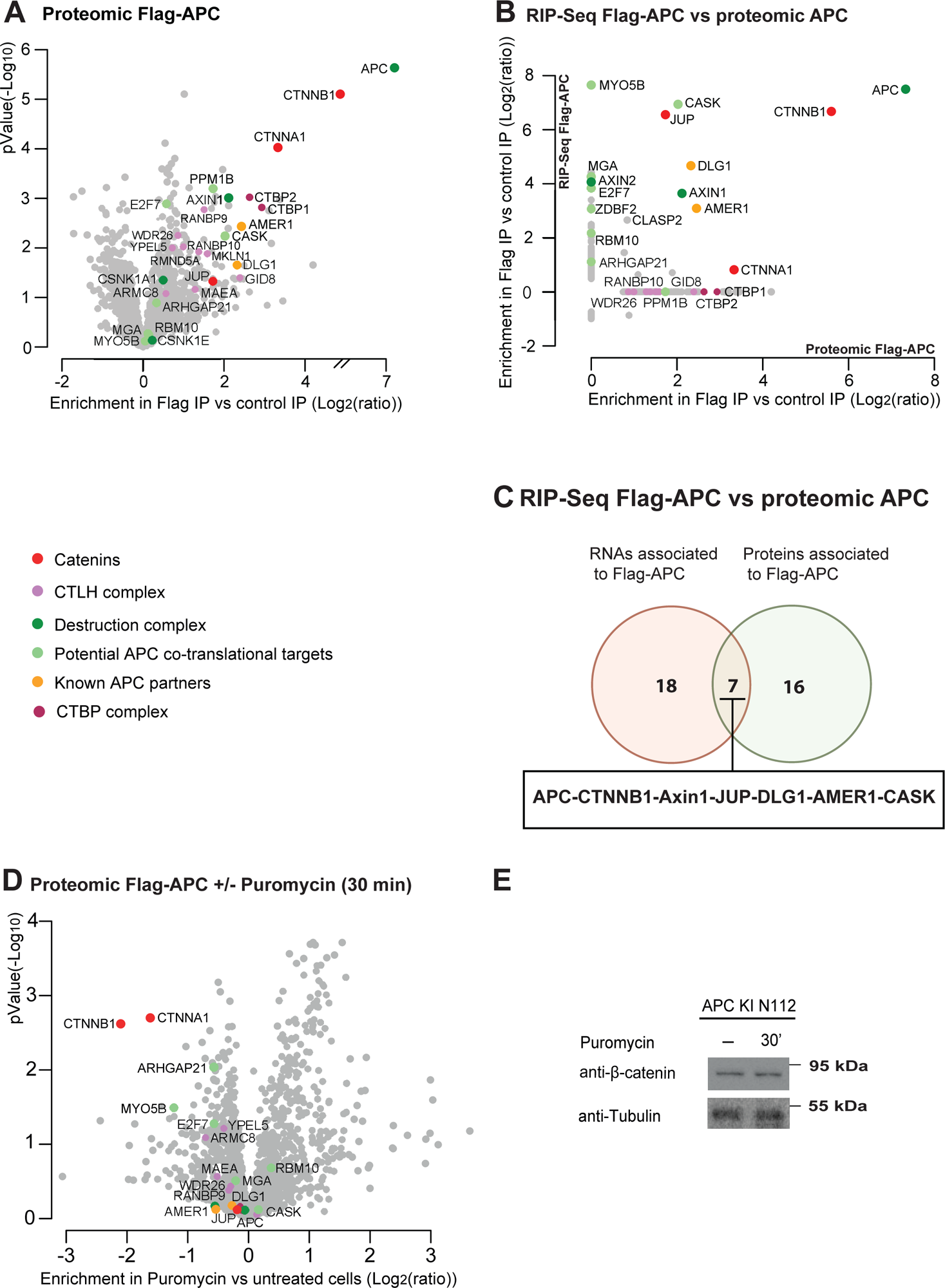
Identification of APC co- and post-translational partners. **A-**Volcano plot representing label-free mass spectrometry analysis of 3xFLAG-mScarlet-APC immuno-precipitates. The x axis displays the enrichment of proteins (log2 [ratio]) immunoprecipitated with anti-FLAG antibody conjugated beads (Flag-IP) of 3xFLAG-mScarlet-APC cells *versus* HEK293 wild-type cells used as control. The y axis represents significance, displayed as -log10 (p-Value). Experiments were done in triplicates and statistical significance is considered for p-Value <0.01. The colored points in the plots represent different categories of proteins, as indicated below the graphs. **B**-The graph shows a comparison of 3xFLAG-mScarlet-APC proteomics *versus* transcriptomics of the first 200 candidates identified in each IP. The x axis represents the enrichment of proteins (log_2_ [ratio]) in FLAG IP *versus* control IP. The y axis displays the enrichment of the RNAs immunoprecipitated in FLAG IP *versus* control IP as log2 [ratio]. **C**-Venn Diagram showing the intersection of the most significant candidates identified with 3xFLAG-mScarlet-APC proteomics (green circle; hits have Log_2_(fold change) ≥ 1.5 and - log_10_(p-Value) ≥ 1) *versus* transcriptomics (red circle; hits have Log_2_fFold change) ≥ 1.1 and - log_10_(p-Value) ≥ 10). **D-**The graph shows a comparison of 3xFLAG-mScarlet-APC interactome before and after translation inhibition with puromycin for 30 min. The x axis displays the enrichment of proteins in puromycin treated *versus* untreated cells as log2 [ratio]. The y axis represents significance, displayed as -log10 (p-Value). Experiments were done in triplicates and statistical significance is considered for p-Value <0.01. **E**-Western blot analysis showing β-catenin levels in 3xFLAG-mScarlet-APC knock-in HEK293 cells, with or without puromycin treatment for 30 min as indicated. Tubulin served as a loading control. Molecular protein weight markers size is indicated on the right. APC KI N112: 3xFLAG-mScarlet-APC knock-in cells, clone N112.

Next, we compared APC transcriptomic *versus* proteomic IPs, and plotted the enrichment values of the FLAG vs. control IP for the first 200 candidates identified in each of the two experiments, as ranked by their p-values (Figure 2B). This showed that APC partners could be grouped in three classes: first, partners that are enriched in the RIP-seq experiment but not in the proteomics, such as MYO5B, MGA, E2F7, ARHGAP21 and RBM10; second, partners enriched in both experiments (β-catenin, Axin1, CASK, DLG1, JUP and AMER1); and finally, partners found significantly enriched only in the proteomics (α-catenin, CTBP1/2, members of the CTLH complex). It is important to note that many partners enriched only in the RIP-seq are still detected in the proteomic experiments, but without enrichment over the control IP (MGA, E2F7, MYO5B, ARHGAP21). Because the interaction of APC with these mRNAs is abolished by puromycin treatment, these results indicate that these proteins represent APC partners that are bound only co-translationally. Conversely, α-catenin binds strictly post-translationally. It is an indirect APC partner that associates via β-catenin (Rubinfeld et al., 1993), possibly explaining why it is not enriched in the RIP-seq experiment while it is a strong APC partner in the proteomics analysis. Using stringent cut-offs value to identify high confidence targets (see legends), these data defined 18 APC partners bound only co-translationally, 7 partners bound co- and post-translationally, and 16 bound only post-translationally (Figure 2C).

### APC co-translational interactions largely contribute to its association with β-catenin

The co-translational interaction of APC with β-catenin might facilitate its recognition by the destruction complex. Indeed, because one APC molecule can bind many β-catenin proteins via repeated motifs present in APC, the binding of APC to a β-catenin polysome would enable it to bind any new nascent protein as soon as they emerge from the ribosome, triggering their degradation. To test this hypothesis, we re-examined the APC protein interactome but this time after removing all co-translational interactions by treating cells with puromycin for 30 minutes. In this case, there is no translation and thus no co-translational interactions. Quantitative mass-spectrometry analysis of the IP’ed proteins showed that APC lost most of its interaction with α- and β-catenin after translation arrest (∼4 fold reduction; Figure 2D, Table S2). This decrease in binding was not due to a change in the expression levels of β-catenin, because Western blots showed that the cellular levels of this protein did not change after puromycin treatment (Figure 2E). Interactions with other proteins were less affected (Figure 2D). These data indicate that the co-translational interactions of APC with β-catenin are required to promote an efficient binding.

### Axin1 associates co-translationally with mRNAs

APC and Axin1/2 interact with each other and form the core of the destruction complex (Ikeda et al., 1998). To further characterize the interaction of the destruction complex with β-catenin mRNA, we performed RNA immunoprecipitation assays using Axin1 as a bait. For this purpose, we used the Flip-in system to stably express a 3xFLAG-eGFP-Axin1 fusion protein. This system is indeed convenient as it allows to express moderate levels of proteins. In HeLa Flip-in cells, 3xFLAG-eGFP-Axin1 was diffusely distributed in the cytoplasm and additionally formed foci as previously described (Figure 3A, Faux et al., 2008; Nong et al., 2021). 3xFLAG-eGFP-Axin1 was immunoprecipitated using anti-FLAG beads while empty beads were used as a control, and the pelleted RNAs were sequenced (Figure 3B, Table S1). Remarkably, Axin1 interacted with β-catenin mRNA, as well as with many other mRNAs including those bound by APC: JUP MYO5B, E2F7, MGA, PPM1B, DLG1, ZDBF2, RBM10 and CASK.

**Figure 3:**
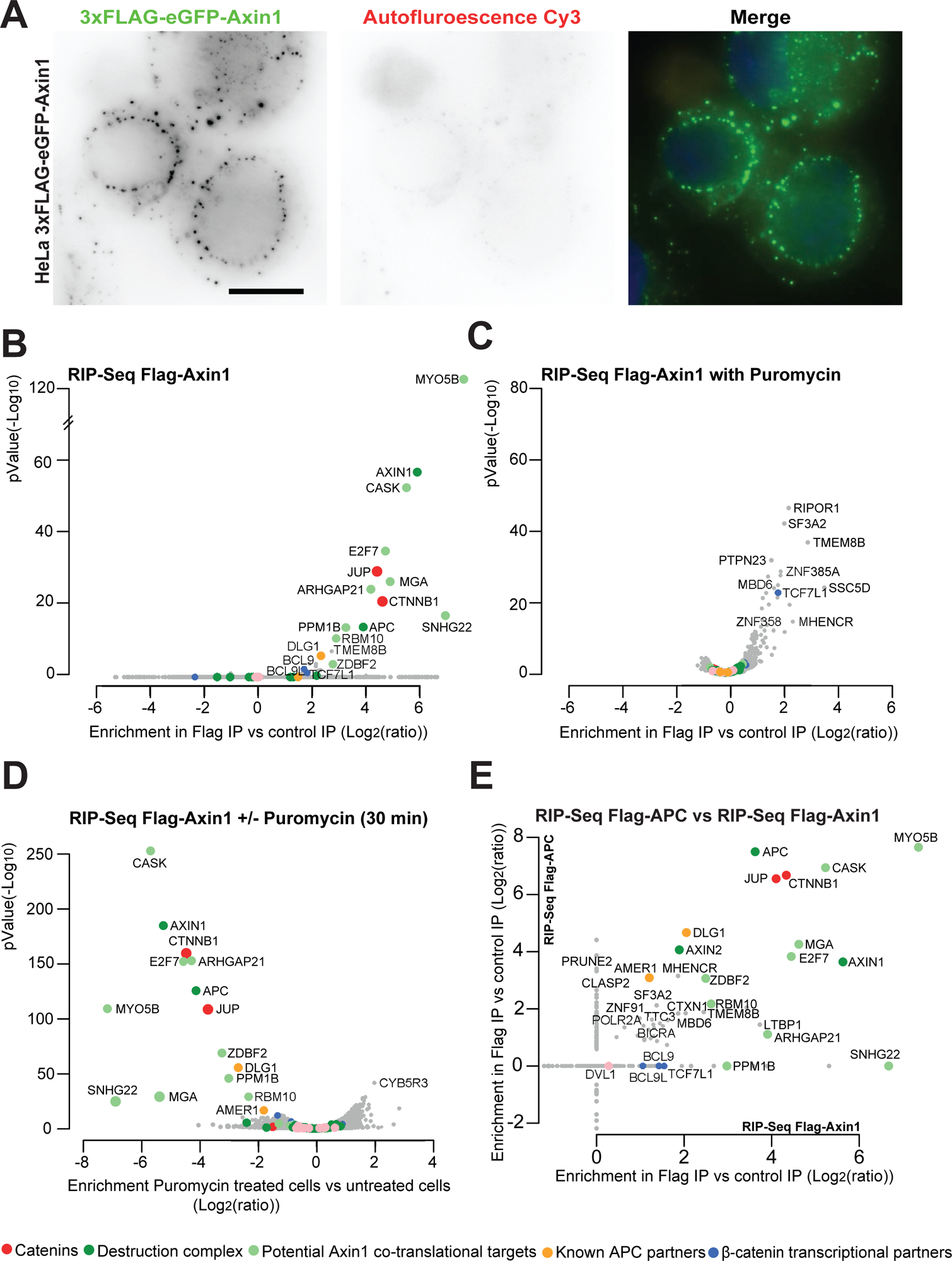
Axin1 interacts with RNA in a translation-dependent manner. **A**-Images are micrographs of HeLa H9 3xFLAG-GFP-Axin1 Flip-In HeLa H9 cells. Left and green: 3xFLAG-GFP-Axin1 signals; middle and red: autofluorescence signals obtained with the DsRed channel. DNA is stained with Dapi. Scale bar: 10 microns **B**-Volcano plots depicting the interaction of 3xFLAG-eGFP-Axin1 with RNAs in HeLa H9 Flip-in cells. The x axis represents the enrichment (log_2_ [ratio]) of the RNAs immunoprecipitated with anti-FLAG antibody conjugated beads (Flag-IP) compared to beads without FLAG antibody (control IP). The y axis represents significance, displayed as -log10 (p-Value). Experiments were done in duplicates and statistical significance is considered for p-Value <0.01. The colored points in the plots represent different categories of proteins encoded by the identified mRNAs. The categories are listed at the bottom. **C**-As in B except that cells were treated with puromycin for 30 min at 100 μg/ml. **D**-Volcano plot comparing RNAs immunoprecipitated in the FLAG IPs, in puromycin *versus* untreated condition. Legend as in C. **E-**The graph shows a comparison of the RNAs enriched in the 3xFLAG-mScarlet-APC IP (HEK293 cells) *versus* the 3xFLAG-eGFP-Axin1 IP (HeLa H9 cells) for the first 350 candidates identified in each IP (ranked by p-Values). The x axis displays the enrichment of the RNAs immunoprecipitated in FLAG IP *versus* control IP (log_2_ [ratio]) in 3xFLAG-eGFP-Axin1 cells. The y axis represents the enrichment of the RNAs immunoprecipitated (log_2_ [ratio]) in FLAG IP *versus* control IP in 3xFLAG-mScarlet-APC cells.

To test whether Axin1 binding to mRNAs was translation-dependent, we repeated the RNA immunoprecipitation after treating cells with puromycin for 30 minutes. As found for APC, the mRNAs that bound Axin1 in normal conditions were not immunoprecipitated after puromycin treatment (Figure 3C-D; S2; Table S1). Interestingly, Axin1 appeared to weakly bind some other mRNAs in puromycin treated cells (Figure 3C). This was the case for the Wnt transcriptional repressor TCF7L1 (TCF3), as well as SF3A2 and RIPOR1, which both appeared to also bind APC in puromycin treated cells (Figure 1D).

Next, we compared the mRNAs bound by APC and Axin1 and plotted the enrichment in the FLAG versus control IP for the top 350 mRNAs of each IP, ranked by p-values (Figure 3E). Many mRNAs associated with both proteins, and this was particularly the case for mRNAs that had high enrichment values over the control IP and high p-values. In fact, the mRNAs that appeared specific for APC or Axin1 had non-significant binding p-values (Figure 3E, Table S1). The exceptions were PPM1B and the non-coding RNA SNHG22, which associated with Axin1 but not APC. Thus, these data show that APC and Axin1 bind co-translationally to a common set of mRNAs, which include β-catenin, MYO5B, CASK, JUP, MGA, E2F7, ARGHAP21, RBM10, ZDBF2 and DLG1 (Figure 3E).

### Axin partners can be strictly co-translational, co- and post-translational, or strictly post-translational

To compare the co-translational partners of Axin1 to its protein interactome, we first IP’ed 3xFlag-GFP-Axin1 and analyzed the pelleted proteins by quantitative mass spectrometry (Figure 4A, Table S2). Our results showed that the expected Axin1 partners were enriched in the Flag IP as compared to the control IP. This included α-β-γ− and δ-catenin (CTNNA1/A2, CTNNB1, JUP, CTNND1), the destruction complex proteins APC, CK1 (CSNK1A1), GSK3α and GSK3β, as well as βTrCP (FBXW11). Other proteins linked to the Wnt pathway were also present, such as AMER1, DLG1, CTBP1/2, as well as other proteins like ARGHAP21 and CASK. Next, we compared the Axin1 partners found in the RIP-seq and proteomic experiments, and for this we plotted the enrichment of the FLAG versus control IP of the top 400 partners, as ranked by their p-values (Figure 4B). As for APC, this defined three types of Axin1 partners: (i) the ones that bind only co-translationally (i.e. enriched in the RIP-seq but not the proteomics), which included MYO5B, MGA, PPM1B, RBM10, E2F7 and ZDBF2; (ii) the partners that bind co- and post-translationally (i.e. enriched in both the RIP-seq and the proteomic IP), such as CTNNB1, APC, ARGHAP21, JUP, CASK, AMER1 and DLG1 (note that except for ARGHAP21 these are the same as for APC; Figure 2); and the partners that bind only post-translationally, such as the kinases of destruction complex (CSNK1A1, GSK3), β-TrCP (FBCW11), α- and δ-catenin, and CTBP1/2. These results are similar to the ones found with APC (Figure 2; Table S1 and S2), indicating that these proteins have similar co-transcriptional and post-transcriptional partners.

**Figure 4:**
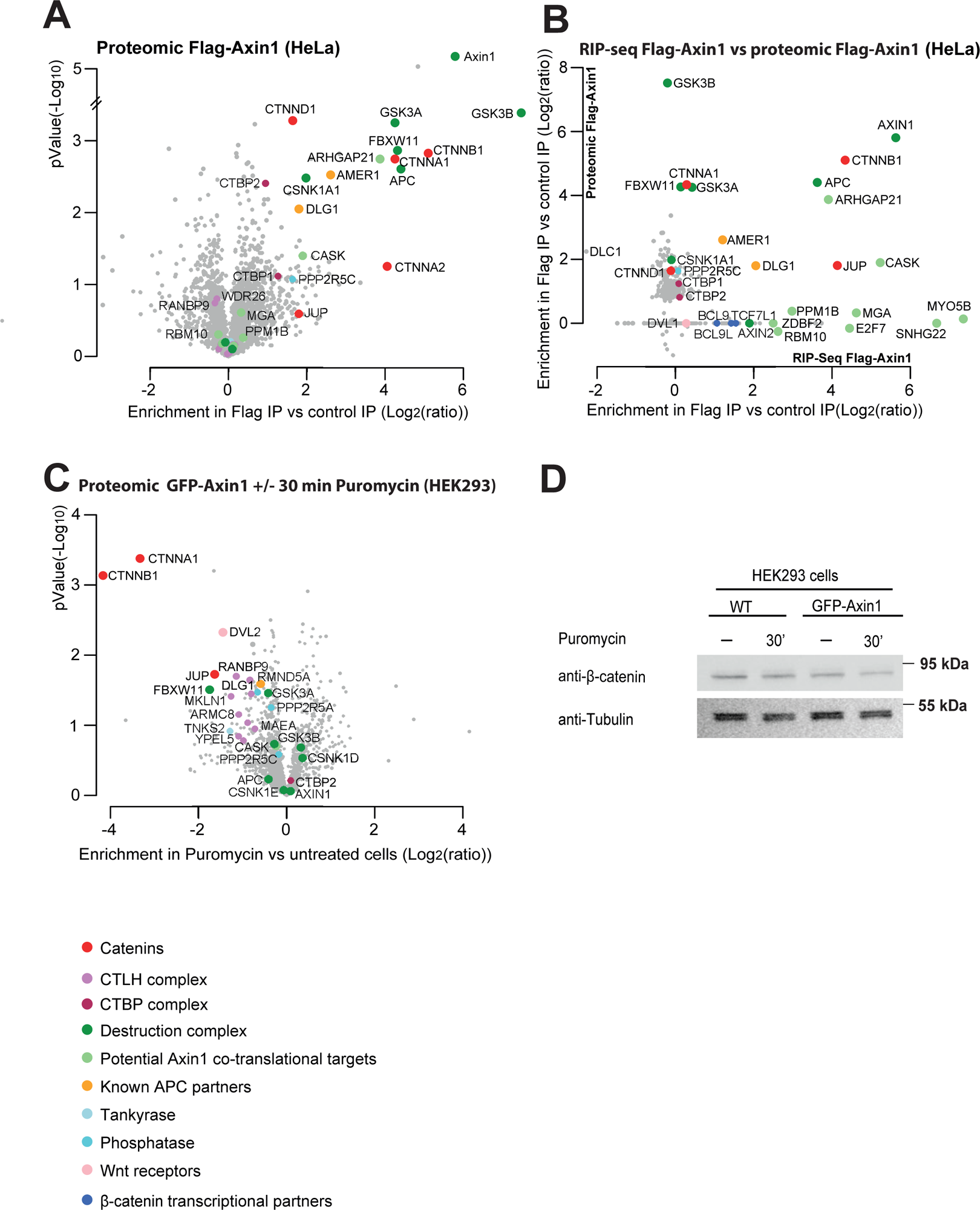
Axin1 interacts predominantly co-translationally with β-catenin. **A-**Volcano plot representing label-free mass spectrometry analysis of 3xFLAG-eGFP-Axin1 immuno-precipitates in HeLa cells. The x axis displays the enrichment of proteins (log2 [ratio]) immunoprecipitated with anti-FLAG antibody conjugated beads (Flag IP) of 3xFLAG-eGFP-Axin1 cells *versus* HeLa H9 wild-type cells used as control. The y axis represents significance, displayed as -log10 (p-Value). Experiments were done in triplicates and statistical significance is considered for p-Value <0.01. The colored points in the plots represent different categories of proteins encoded by the identified mRNAs. **B**-The graph shows a comparison of 3xFLAG-eGFP-Axin1 proteomics *versus* transcriptomics for the first 400 candidates identified in each (HeLa H9 cells). The x axis displays the enrichment of the RNAs immunoprecipitated in FLAG IP *versus* control IP as log2 [ratio]. The y axis represents the enrichment of proteins (log_2_ [ratio]) in FLAG IP *versus* control IP. **C-**Volcano plot comparing proteins immunoprecipitated in eGFP-Axin1 IPs in HEK293 cells, in puromycin *versus* untreated condition. The x axis displays the enrichment of proteins immunoprecipitated in GFP IPs, in puromycin treated *versus* untreated cells (Log_2_ [ratio]). The y axis represents significance, displayed as -log_10_ (p-Value). **D**-Western blot analysis showing β-catenin levels in wild-type and eGFP-Axin1 expressing HEK293 cells, with or without puromycin treatment for 30 min as indicated. Tubulin served as a loading control. Molecular protein weight markers size is indicated on the right. (-): untreated condition; (30’): 30 min puromycin treatment; WT: wild-type HEK293 cells; GFP-Axin1: HEK293 cells expressing eGFP-Axin1.

## Axin1 binding to β-catenin is mainly co-translational

Next, we tested the role of the co-translational interaction of Axin1 and compared its protein interactome with and without co-translational interactions, by briefly inhibiting translation with puromycin. As in the case of APC, 30 minutes of puromycin treatment led to a ∼10 fold decrease in binding to α- and β-catenin (Figure S3A). Because we found that β-catenin levels diminished in HeLa cells treated with puromycin (Figure S3B), we repeated this experiment in HEK293 cells. Indeed, β-catenin protein expression levels in these cells were unaffected by the puromycin treatment (Figure 4D). We thus established a HEK293 cell line stably expressing GFP-Axin1, and performed IP assays using anti-GFP beads and empty beads as control, with and without a 30 minutes puromycin treatment. In untreated conditions, many expected Axin1 partners were enriched in the GFP versus control IP (Figure S3C, Table S2), and the results were similar to the ones of HeLa cells (Figure 4). Among the few differences, the phosphatase PP2A and the CTLH ubiquitin ligase complex were pulled-down in HEK293 but not Hela cells, while CTNND1, AMER1, CTBP1/2 and CASK were more enriched in the HeLa IPs.

Next, we compared Axin1 partners in puromycin versus untreated conditions (Figure 4, Table S2). Again, the interaction of Axin1 with α- and β-catenin decreased 10-16 fold after a 30 minute puromycin treatment (Figure 4C). The other proteins most affected by the arrest of translation were βTrCP (FBXW11), which may bind Axin1 less tightly in absence of β-catenin (Su et al, 2008) and γ-catenin (JUP), the β-catenin paralog. These effects were specific because binding of Axin1 to the other members of the destruction complex was little or not affected. The strong decrease of the association of Axin1 to β-catenin upon translation arrest indicates that the co-translational interactions constitute a large fraction of the total interactions of Axin1 with β-catenin.

### APC regulates the expression of some of its co-translational partners

APC induces the degradation of β-catenin. As shown in Figure 1C and 2C, we detected several co-translational partners of APC in addition to β-catenin, suggesting that APC may also regulate their expression. To address this, we depleted APC with siRNA and assessed protein levels and localization using Western blots and immunofluorescence (IF). In these experiments, we used the β-catenin BAC cell line and monitored β-catenin and β-catenin-GFP as positive controls (Figure 5). As expected, β-catenin levels increased ∼6.5 fold after APC depletion, while a control FFL siRNA had no effect (Figure 5A). We tested DLG1, CASK, ARGHAP21 and JUP (γ-catenin), as examples of APC co-translational targets. We observed that knocking down APC increased expression of JUP by 1.9 fold and had more modest effects on CASK and ARGHAP21 (1.3 and 1.4 fold increase; Figure 5A). No effects were observed on the levels of DLG1. Next, we analyzed protein localization by immunofluorescence (IF). β-catenin concentrated at cell-cell junctions in cells treated with the control siRNA while it accumulated in the nucleus after APC know-down, as expected (Figure 5B). JUP (γ-catenin), the β-catenin paralog that is regulated similarly to β-catenin (Kodama et al., 1999; Rubinfeld et al., 1995), also localized at cell-cell junction in control cells and relocalized to the nucleus after APC knock-down (Figure 5C). Interestingly, we also observed a change in the localization of ARHGAP21. In control conditions, this protein localized in puncta at the periphery of cell, as previously described (Sousa et al., 2005), but it showed a diffuse cytoplasmic signal after APC knock-down (Figure 5D). Taken together, these results show that APC downregulates the levels of some of its co-translational targets, namely JUP and CASK, and that its removal leads to the intracellular relocalization of JUP of ARGHAP21, as is the case for β-catenin.

**Figure 5:**
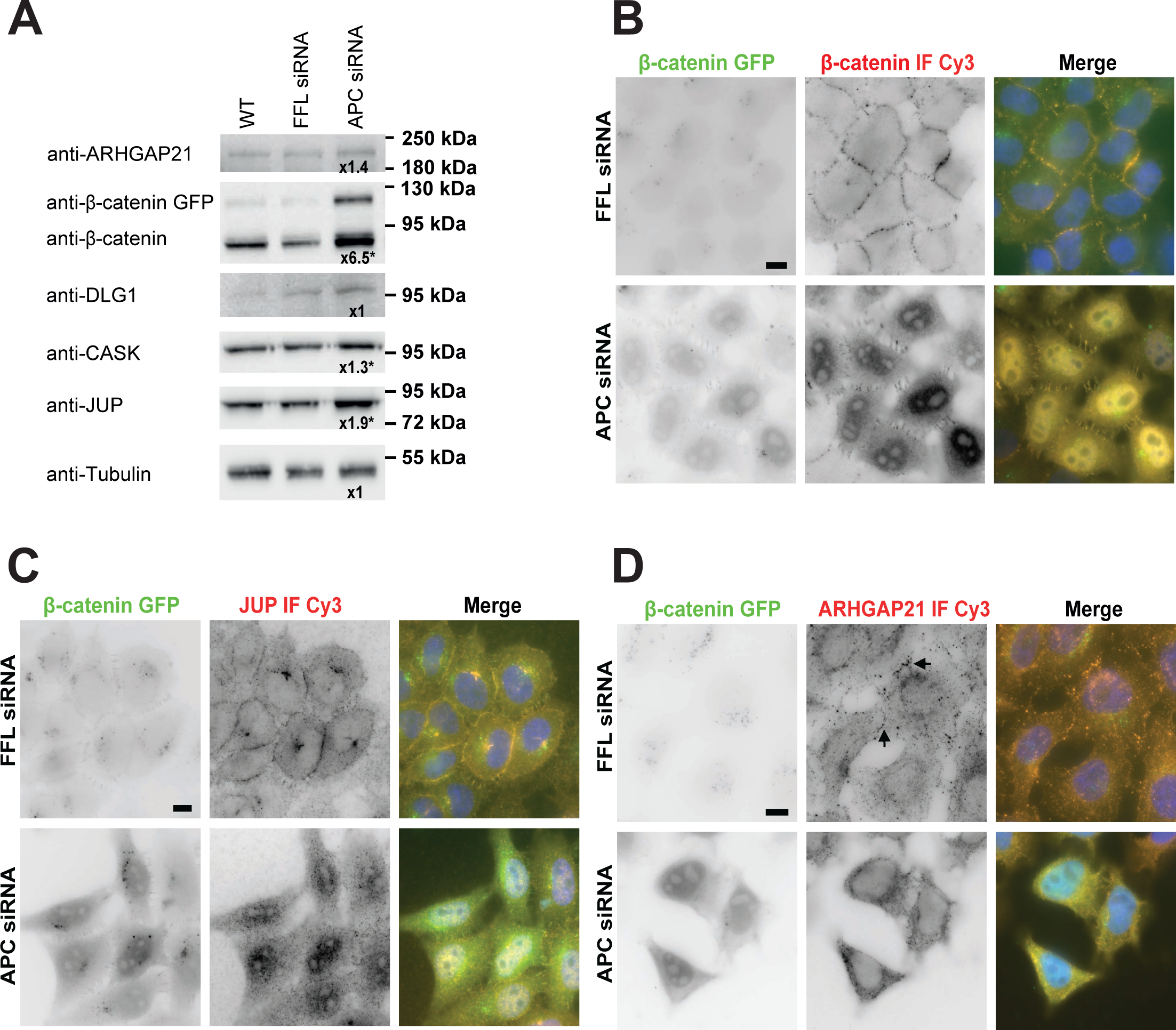
Effects of APC depletion effects on the expression and localization of its co-translational targets. **A-**Western blot analysis showing proteins levels in HeLa β-catenin-GFP BAC cells before and after APC knock-down with siRNAs. Cells were treated with an APC siRNA or a control firefly (FFL) siRNA as indicated. Tubulin served as a loading control. Immunoblotted proteins are displayed on the left. Molecular protein weight markers size is indicated on the right. WT: untreated cells. Protein expression variations are ratios of levels in APC siRNA vs FFL siRNA conditions, after normalization with tubulin, and are indicated under each blot. β-catenin levels were calculated as a mean of both endogenous β-catenin and exogenous β-catenin GFP levels. Experiments were done in triplicates and p-Value <0.05 with a unilateral paired t-test are indicated by stars. **B-**Images are micrographs of HeLa β-catenin-GFP BAC cells treated with control FFL siRNA (upper) or with APC siRNA (lower). Left and green: signals corresponding to β-catenin-GFP; middle and red: Cy3 fluorescent signals corresponding to β-catenin protein detected with Immunofluorescence. DNA is stained with DAPI. Scale bar: 10 microns. **C-**As in B except that middle and red panel represents Cy3 immuno-fluorescent signals of JUP antibodies. **D-**As in B except that middle and red panel represents Cy3 immuno-fluorescent signals of ARHGAP21 antibodies.

### The β-catenin mRNA condensates do not contain other APC-bound mRNAs

In absence of Wnt, the β-catenin mRNAs form foci that contain multiple mRNA molecules as well as APC and Axin (Chouaib et al., 2020; Figure 1A). These foci likely form via the condensation of β-catenin polysomes induced by the multivalent interactions of APC and Axin with nascent β-catenin protein (Chouaib et al., 2020; see below). It was thus interesting to see whether the other APC bound mRNAs would also accumulate in the β-catenin mRNA condensates. To test this possibility, we used the β-catenin-GFP BAC cell line and performed two-color smFISH as well as smFISH in combination with APC IF (Figure 6 and Figure S4). The APC staining showed large foci that colocalized with the β-catenin mRNA condensates as well as tiny spots (see Figure 1A above). Importantly, the APC tiny spots occasionally colocalized with single β-catenin mRNAs, suggesting that the antibody was sensitive enough to detect interaction with single polysomes (see Figure 1A above, orange arrows). In contrast to β-catenin mRNAs, JUP mRNAs did not form condensates and were present as dispersed single molecules (Figure 6A and B). In addition, this mRNA did not accumulate in the β-catenin mRNA condensates, identified either by β-catenin smFISH or by the large APC foci that were used as proxy (Figure 6A and B). Moreover, no particular co-localization could also be observed between JUP and β-catenin mRNAs outside foci. Interestingly however, some single molecules of JUP mRNAs colocalized with APC spots (Figure 6A, orange arrows), consistent with the co-translation interactions observed by RIP-seq (see above). Very similar results were obtained for CASK, MYO5B, DLG1, AMER1 and ARHGAP21 mRNAs, which neither formed large condensates nor accumulated in β-catenin mRNA foci, but occasionally colocalized with tiny APC spots (Figure 6A-B and S4A-B).

**Figure 6:**
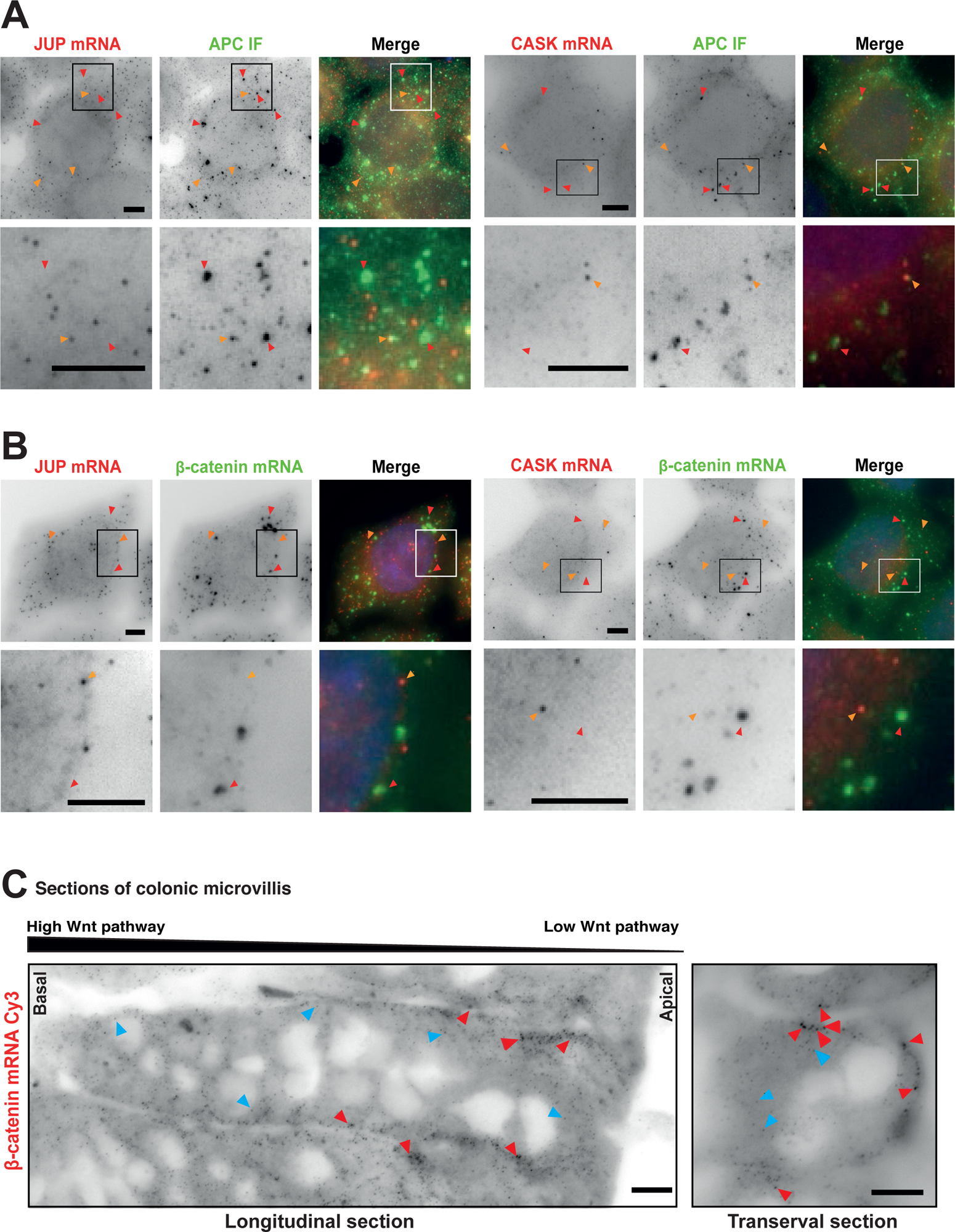
β-catenin mRNAs form homotypic foci. **A-**Images are micrographs of HeLa β-catenin GFP BAC cells. Left and red: Cy3 fluorescent signals corresponding to JUP or CASK mRNAs detected with smiFISH; middle and green: FITC signals corresponding to APC detected by immunofluorescence. For each cell, a zoom of the boxed shown below. Red arrows show APC foci and orange arrows show single mRNAs of JUP or CASK. DNA is stained with DAPI. Scale bar: 5 microns. **B-**Images are micrographs of HeLa β-catenin GFP BAC cells. Left and: Cy3 signals corresponding to JUP or CASK mRNAs detected with smiFISH; middle green: Cy5 fluorescent signals corresponding to β-catenin mRNAs detected with smFISH. For each cell, a zoom of the boxed area is shown below. Red arrows show β-catenin mRNA foci and orange arrows show single mRNAs of JUP or CASK. DNA is stained with DAPI. Scale bar: 5 microns. **C-**Images are micrographs of mice wild-type colonic tissue with a crypt longitudinal section (left) and transversal section (right). Panels represent Cy3 fluorescent signals corresponding to β-catenin mRNA detected with smiFISH. Red arrows show β-catenin foci and blue arrows show β-catenin mRNA single molecules. DNA is stained with DAPI. Scale bar: 10 microns.

β-catenin mRNA condensates have been observed in the β-catenin-GFP BAC cell line and in HEK293 cells (Chouaib et al., 2020; Figure 1A), but it is currently unknown whether normal tissues also have these mRNA foci. We thus generated frozen sections of mouse colons and performed smFISH on longitudinal and transversal crypt sections, which have an oriented Wnt gradient as indicated in Figure 6C. The results showed that intestinal epithelial cells had β-catenin mRNAs occurring as both single molecules and foci (Figure 6C and S5A). Taken together, these data indicate that β-catenin mRNA condensates are physiologically relevant structures that are homogeneous in terms of RNA content.

### β-catenin mRNAs form two polysome populations that interact with either APC or E-cadherin

The previous results show that β-catenin is co-translationally bound by APC and Axin1, suggesting that little or no newly made β-catenin escapes degradation in absence of Wnt. β-catenin is however a bi-functional protein that also plays a key role at cell-cell junctions, where it links E-cadherin to the actin cytoskeleton (Drees et al., 2005). The co-translational interactions of β-catenin with APC and Axin thus question the source of the β-catenin localizing at the cell membrane. One possibility that could solve this conundrum would be that there are in fact two populations of β-catenin polysomes, one bound by APC and Axin and functioning in the Wnt pathway, and another one specialized in producing plasma membrane β-catenin. A simple way to generate this second type of polysomes would be to have them interact with E-cadherin instead of APC and Axin. To test this hypothesis, we stably expressed an E-cadherin-GFP fusion in HEK293 cells and analyzed the associated mRNAs by RIP-seq. E-cadherin-GFP protein localized to the cell membrane as expected (Figure S6A). We performed an IP using anti-GFP beads, with and without a 30 minutes puromycin treatment. The results show that in untreated conditions, E-cadherin-GFP interacted with only three mRNAs (Figure 7A, Table S1): β-catenin, JUP (γ-catenin) and GOLGA4 mRNAs, which encodes a protein of the Golgi apparatus involved in E-cadherin trafficking (Lock et al., 2005). Puromycin treatment abolished these interactions, indicating that they are indeed co-translational and mediated by the nascent protein chains (Figure 7B, Table S1). These results suggest that the interactions between E-cadherin and β-catenin mRNA could occur at the cell membrane. Indeed, smFISH showed that some single molecules of β-catenin mRNA localize at the cell membrane where they overlapped with the E-cadherin-GFP signal (Figure S6B). Moreover, a three color experiment performed on β-catenin BAC cells expressing E-cadherin GFP, showed that β-catenin mRNAs were localized either at the cell membrane with E-cadherin or in the cytoplasm with APC (as single mRNAs or mRNA condensates), but not with both at the same time (Figure 7C). These results were consistent with the idea that β-catenin mRNAs form two distinct populations of polysomes, one that interacts with APC and Axin, mainly in condensates, and makes a protein destined for degradation, and one that interacts with E-cadherin at the plasma membrane and synthesizes proteins functioning in cell-cell adhesion. To further confirm this model, we analyzed the E-cadherin-GFP interactome by label-free quantitative proteomic. E-cadherin-GFP interacted with catenins (α, β and δ) and other known partners, but no interaction was found with APC or Axin, as no peptides for these proteins were detected (Figure 7D, Table S2). Conversely, in the APC and Axin proteomics, no E-cadherin peptides were detected (Figure 2A-4A-S3C; see Table S2). Taken together, these data provide solid evidence that E-cadherin interacts co-translationally with some β-catenin polysomes and transport these molecules to the plasma membrane, away from the destruction complex. APC and Axin interact with other β-catenin polysomes and condense them in cytoplasmic foci, where they are unlikely to interact with E-cadherin. The sorting of β-catenin mRNAs to two different intracellular locations is likely reinforced by the mutually exclusive interactions of APC and E-cadherin for β-catenin protein (Hiilsken et al., 1994).

**Figure 7:**
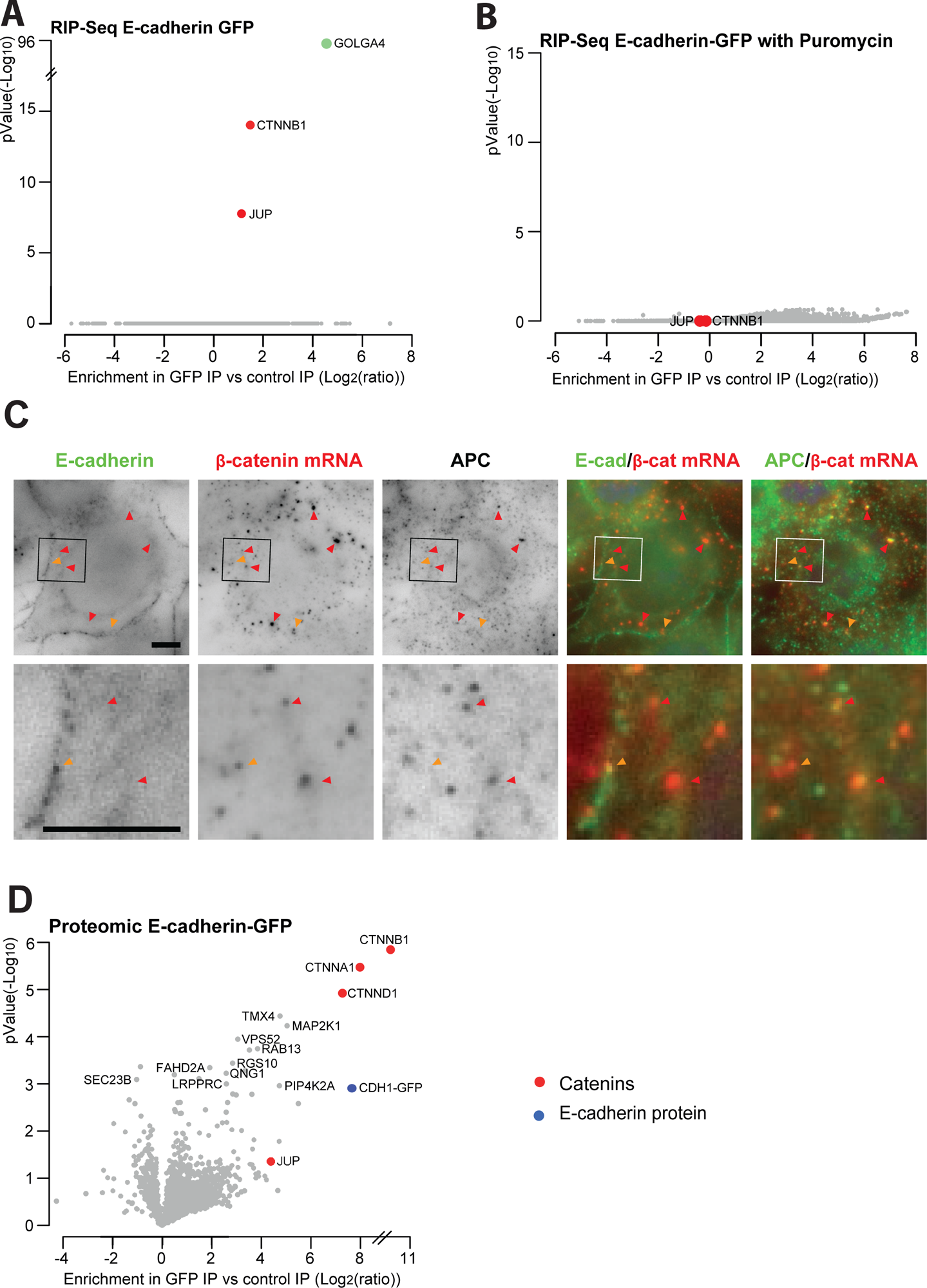
E-cadherin interacts with β-catenin in a translation-dependent manner. **A**-Volcano plots depicting the interaction of E-cadherin-eGFP with RNAs in HEK293 cells. The x axis represents the enrichment (log_2_ [ratio]) of the RNAs immunoprecipitated with anti-GFP antibody conjugated beads (GFP-IP) compared to beads without GFP antibody (control IP). The y axis represents significance, displayed as -log10 (p-Value). Experiments were done in duplicates and statistical significance is considered for p-Value <0.01. The colored points in the plots represent different categories of proteins encoded by the identified mRNAs. **B**-As in A except that cells in the GFP and control IPs were treated with puromycin for 30 min at 100 μg/ml. Legend as in A. **C-**Images are micrographs of HeLa BAC cells expressing E-cadherin-eGFP. First left panel: E-cadherin-eGFP signals; second panel: Cy3 fluorescent signals corresponding to β-catenin mRNAs detected with smiFISH; third panel: Cy5 fluorescent signal corresponding to APC protein detected with immunofluorescence. Color overlays are shown two channels at a time with the indicated false colors. For each cell, a zoom of the boxed area is shown below. Orange arrows show β-catenin mRNA co-localizing with E-cadherin and red arrows show β-catenin mRNAs co-localizing with APC foci. DNA is stained with DAPI. Scale bar: 5 microns. **D-**Volcano plot representing label-free mass spectrometry analysis of E-cadherin-GFP immuno-precipitates. The x axis displays the enrichment of proteins (log2 [ratio]) immunoprecipitated with anti-GFP antibody conjugated beads (GFP-IP) of E-cadherin-GFP cells *versus* HEK293 wild-type cells used as control. The y axis represents significance, displayed as -log10 (p-Value). Experiments were done in triplicates and statistical significance is considered for p-Value <0.01.

## Discussion

This study reveals that APC and Axin bind co-translationally to a number of mRNAs, that the interactions of the β-catenin protein with APC and Axin are predominantly co-translational and that the β-catenin mRNA generates two types of specialized polysomes, coding for the same protein but with different partners and functions.

### Co-translational interactions sort β-catenin polysomes to different locations and functions

A key result of this work is the discovery that β-catenin mRNA makes co-translational interactions with APC and Axin on one side, and E-cadherin on the other. In the cell, β-catenin mRNAs co-localize with either E-cadherin at the plasma membrane, or with APC in the cytoplasm, in the latter case as single molecule or polysome condensates. Proteomic analysis of E-cadherin, APC and Axin further show that while all three proteins associate with β-catenin, E-cadherin does not show any interaction with APC and Axin, and conversely, APC and Axin fail to pull down E-cadherin (Table S2). This is consistent with the idea that the known mutually exclusive interactions of APC and E-cadherin for β-catenin (Hiilsken et al., 1994) take place at the level of entire polysomes. This way, the co-translational interactions of β-catenin creates spatially separated polysome populations that are specialized for one of the functions of this multifunctional protein, a mechanism that we termed polysome sorting. When a β-catenin mRNA reaches the cytoplasm and start being translated, some of the nascent β-catenin protein may bind the destruction complex. Due to the ability of APC to bind numerous β-catenin proteins with repeated motifs (Rubinfeld et al., 1993), all subsequent polypeptides translated by this polysome will have a high probability of also binding the destruction complex. Indeed, APC could move over nascent β-catenin proteins being elongated, much like a person exercising on a treadmill and walking while staying at the same place. APC could thus remain stably bound to a single β-catenin polysome despite the very dynamic nature of both translation and the interactions involved. Interaction of APC with β-catenin polysomes may be further reinforced if the mRNA joins the polysome condensates, which have a high concentration of APC and Axin (Chouaib et al., 2020, this work). In contrast, if the nascent protein of a freshly exported β-catenin mRNA binds E-cadherin, the polysome will become associated to the plasma membrane or to trafficking vesicles, and it will remain away from the destruction complex, also favoring interactions of subsequently synthesized β-catenin with neighboring E-cadherin molecules. Given the translation elongation rate (5-10 amino acids per second, or ∼2.5 min for the entire β-catenin protein; Gerashchenko et al., 2021), and the kinetics of ER to plasma membrane transport (tens of minutes; Hirschberg et al., 1998), it is unlikely that E-cadherin can transport a β-catenin polysome from the ER to the plasma membrane, unless ribosome pause for a long time. Rather, it appears more likely that β-catenin polysomes bind E-cadherin proteins that are either at the plasma membrane or cycling between the plasma membrane and endosomes. Since E-cadherin molecules not engaged at adherens junctions are constantly cycling through the endosomal system (Lock et al., 2005), this may provide a mechanism to transport β-catenin mRNAs to the plasma membrane as these E-cadherin molecules are precisely the ones lacking β-catenin. This raises the question of how the polysome sorting mechanism is regulated. The affinities of β-catenin for APC/Axin and E-cadherin are positively regulated by phosphorylation (Ha et al, 2004), and E-cadherin phosphorylation occurs early before the protein reaches the plasma membrane (McEwen et al., 2014), while APC/Axin phosphorylation is regulated by Wnt signaling (Pronobis et al., 2015; Yamamoto et al., 1999). Biochemical data indicate that the affinity of β-catenin for phosphorylated E-cadherin is exceptionally high (∼50 pM), 2-3 orders of magnitude higher than the interaction for phosphorylated APC (Choi et al., 2006). This suggests a model in which E-cadherin binds first to β-catenin polysomes and sort these polysomes to the plasma membrane until all E-cadherin molecules are saturated. Consistently, only a minority of β-catenin polysomes are located at the plasma membrane at steady state (see Figure 7C and S6B), when most E-cadherin molecules are expected to be already associated with β-catenin. The remaining β-catenin polysomes would then be available for binding APC and Axin.

Polysome sorting likely applies to other mRNAs. JUP encodes a β-catenin paralog (γ-catenin) that localizes to adherens junctions and accumulates in the nucleus in a Wnt/APC dependent manner (Kolligs et al., 2000; this study). JUP mRNA interacts co-translationally with APC/Axin and E-cadherin, suggesting that like β-catenin mRNA, it is able to form two distinct populations of polysomes. ARHGAP21 binds co-translationally to APC and Axin, and while the protein normally localizes in puncta at the cell periphery, it becomes diffusely distributed in the cytoplasm upon APC knock-down (Figure 5D). This suggests that APC may regulate a different pool of ARHGAP21 than the one making the peripheral protein. More generally, the co-translational targets of APC identified here constitute potential candidates for polysome sorting.

### APC and Axin as co-translational regulators

It is striking that a short puromycin treatment nearly abolishes the interactions of APC and Axin with the β-catenin protein. This means that at steady-state, most of the interactions take place co-translationally, directly on β-catenin mRNAs. APC and Axin can thus be viewed as co-translational regulators, an idea that is reinforced by the fact that the interaction of APC and Axin with a number of the partners identified in this study is detected only at the co-translational level. These proteins may thus form a class of regulators specifically acting co-translationally. In the case of APC and Axin1, this function may be related to or independent of their roles in the destruction complex. An interesting possibility would be that APC and Axin1 can affect the translation dynamics of β-catenin as well as that of the other identified proteins. For instance, β-catenin ribosomes may pause in absence of co-translational binding, waiting to join APC, Axin or E-cadherin. On the contrary, binding of APC and Axin may lead to an arrest of elongation, triggering ribosome collisions and their removal.

### APC, a non-canonical RNA-binding protein or an RNA-proximal protein

Previous work has shown that APC can be cross-linked to mRNAs in vivo, including to that of β-catenin (Preitner et al., 2014; Trendel et al., 2019). However, APC does not have a canonical RNA binding domain, which may suggest that it is as an atypical RNA binding protein (Preitner et al., 2014). Our study confirms the binding of APC to mRNAs but it also demonstrates that this occurs via the nascent protein rather than directly to the mRNA. Thus, the cross-links observed previously with β-catenin mRNA are likely due to random collisions induced by physical proximity. Over the last ten years, it has been shown that a large number of proteins can cross-link to mRNAs, including a large category that do not have an RNA binding domain and are cataloged as ‘atypical RNA binding proteins’ (Albihlal & Gerber, 2018; Hentze et al., 2018). Our data suggest that like APC and Axin1, some of these proteins may cross-link to RNA via a proximity effect rather than by direct and specific binding. We propose to refer to these proteins as ‘RNA proximal proteins’ in order to illustrate their physical proximity to RNAs.

### APC and Axin specifically condensate β-catenin polysomes

In absence of a Wnt signal, β-catenin mRNAs form cytoplasmic foci containing APC and Axin1, and knock-down of these proteins dissolve the β-catenin mRNA condensates (Chouaib et al., 2020). Given the multivalency of APC and Axin1 for themselves and β-catenin (Rubinfeld et al., 1993), and the fact that β-catenin polysome contain multiple β-catenin nascent chains, the co-translational interactions found here provide solid evidence in favor of a model in which APC and Axin1 bridge different β-catenin polysomes to drive their condensation. Surprisingly, we observe that the other mRNAs bound co-translationally by APC and Axin1 are not part of the β-catenin polysome foci. These are thus homotypic condensates, as previously described for germ granules in Drosophila (Trcek et al., 2020). The absence of other mRNAs in the β-catenin condensates could be due to a lower multivalency of the system, caused for instance by a low number of ribosomes on these mRNAs or a monovalent binding of APC to these proteins. It could also be due to a yet unknown property of the β-catenin system, which may enable it to recruit additional partners favoring polysome condensation. In any case, we observe that the β-catenin mRNA foci occur in normal mouse colonic tissue, indicating that they have physiological importance. Future studies will clarify their roles in β-catenin regulation.

## Acknowledgments

We thank Hussein Karaki for his contribution in cell sorting experiments. We thank the facilities MRI, PPM and MGX for their participation in both experiments and analysis. Mass spectrometry experiments were carried out using the facilities of the Montpellier Proteomics Platform (PPM) and RNA sequencing experiments were performed by Montpellier GenomiX Plateform (MGX). MGX acknowledges financial support from France Génomique National infrastructure, funded as part of ‘Investissement d’Avenir’ program managed by Agence Nationale pour la Recherche (contract ANR-10-INBS-09). We acknowledge support from the National Infrastructure France-BioImaging supported by the French National Research Agency (ANR-10-INBS-04). We thank La ligue contre le Cancer (LNCC) and l’Association pour la Recherche sur le Cancer (ARC) for their fellowships to SS. The projet was funded by MSDAvenir grant RNAcan!, and by the Agence Nationale de la Recherche (grant ANR-19-CE12-0007) and the Institut National de cancer (grant PLBIO21-222).

## Author contributions

The project was conceived by E.B. and developped by S.S., who did all the experiments. M.S. and K.E.K performed the MS analysis and analzed these data. S.R. and S.G. performed the sequencing and analyzed these data. The mouse work was done by B.L., M.H. and S.S. Data interpretation was performed by C.E., M.H., S.S. and E.B. The two polysome concept was developped by S.S., C.E. and E.B. The manuscript was written by S.S. and E.B., reviewed and edited by all authors.

## STAR Methods

### Cell culture

HeLa and HEK293 cells were grown in Dulbecco’s modified Eagle’s Medium DMEM (Gibco #31966-021) supplemented with 10% fetal bovine serum (Sigma Aldrich) and 1/100 U/ml Penicillin/streptomycin (Gibco #15140-122). All cells were grown at 37°C with 5% CO2. HeLa β-catenin GFP BAC cells (cells #3843, a gift from T. Hyman lab) were cultured with 400 μg/ml G418. For clone selection in HEK293 cells, puromycin was used at 1 μg/ml and G418 at 300 μg/ml. In HeLa H9 Flp-In cells, hygromycin was used at 50 μg/ml. For translation inhibition assays, puromycin was added to cells at 100 μg/ml for 30 min. For Axin1 stabilization, LG007 (Selleckchem #S7239) was used for 2h at 0.5 nM. All treatments were done at 37°C. Transfection was performed using JetPrime (Polyplus #114-75), at around 50% confluence, 1 day after seeding the plates.

### Generation of CRISPR Cas9 knock-in cells

To tag APC at its N-terminus, the repair cassette for the APC gene contained two 500 bp homology arms flanking the puromycin resistance gene, a P2A sequence, a 3xFLAG tag and the red fluorescent protein the mScarlet (Figure SA). The guides sequence used for the APC knock-in were as follow: APC gRNA1 5’ CACCGTGGCAAGATCTGAGGGAGTT 3’; APC gRNA2 5’ CACCGAGAAAATCAGGATACCCTAC 3’. The puc57-attbU6-sgRNA vector was based on Chen et al. (Chen et al., 2015). The guides RNAs were cloned into this expression vector using BbsI restriction enzyme. Plasmids were verified by sequencing. Cas9-nickase expressing plasmid was provided from Addgene (#42335). The PAM and/or the guide binding sites were mutated in the repair cassettes. Plasmids were transfected in a 3:3:1 weight ratio (repair cassette, each U6 guide, Cas9 Nickase) using JetPrime. Cells were split after 24h and put under selection. Two to three weeks after transfection, individual clones were picked using cloning discs (Merck #F37847-0001) and cultured in 24 well plate for several weeks under puromycin selection. Clones were then verified by genotyping, fluorescent microscopy, and Western blot.

### Generation of stable cell lines using the Flip-In system

The coding sequence of Axin1 was fused at the C-terminus of a 3xFLAG-GFP tag in a pcDNA5 vector, under the control of a CMV promoter. This construct was transfected with a plasmid expressing the flippase protein (Addgene; #13793) in HeLa H9 cells using 1:2 ratio, using JetPrime. Cells were split 24h after transfection and clones were selected in hygromycin. Individual clones were picked, expanded and checked using fluorescent microscopy and Western blot.

### Generation of stable cell lines using retroviral infection

#### Virus production

HEK 293T cells were plated in a 10 cm plate and transfected overnight with a mixture containing 8 μg of the lentiviral plasmid, alongside various plasmids coding for viral accessory proteins (0.4 μg of GAG, POL and TAT/REV plasmids and 0.8 μg of VSV-G). Medium containing viral particles was collected each day for 2 days. Viral particles were concentrated using LentiX concentrator (TaKaRa bio #631232) according to the manufacturer’s instructions then centrifuged at 1500 g for 1 hour at 4°C. Virus particles were suspended in serum-free medium and aliquoted for single use. Aliquots were stored at -80°C for several weeks.

#### HEK293 GFP-Axin1 and E-cadherin-GFP cells

Wild-type HEK293 cells were infected with viruses expressing GFP-Axin1 (N-terminal tag) or E-cadherin-GFP (C-terminal tag). Pools of lowly expressing, GFP positives cells were sorted using FACS. GFP expression and proteins localization were then assessed using fluorescent microscopy and Western Blots.

#### β-catenin BAC E-cadherin-GFP cells

β-catenin BAC cells were infected with viruses expressing E-cadherin-GFP (C-terminally tag). GFP expression and protein localization were assessed using fluorescent microscopy and Western blots.

### PCR Genotyping

Genomic DNAs were prepared using the Promega kit #A1125. PCR amplifications were performed using Taq platinum polymerase (Promega #M780B) or Phusion polymerase (NEB #M0530L). APC CRISPR knock-in clones were verified by PCR using the following primers: 5’ GTATCTTTTGGCCTCAGTTTGAG and 5’ GTGGGCTTGTACTCGGTG, for the 5’ homology arm; 5’ CCACCTACAAGGCCAAGAAG and 5’ CGATTTTCTAACCTGTCTTGGAGG for the 3’ homology arm. PCR products were sequenced to verify proper recombination.

### Western blotting

Cells were lysed using HNTG lysis buffer (20 mM Hepes (pH=7,9), 150 mM NaCl, 1% triton, 10% glycerol, 1 mM MgCl_2_ and 1 mM EGTA) supplemented with 1x proteinase inhibitor cocktail (Roche #54925800). Total proteins concentration was measured using BCA Kit (Thermo-scientific #23227) according to the manufacturer’s instructions. 20 μg to 60 μg of proteins were loaded in 10% SDS page for proteins < 150 kDa, and 8% SDS page for proteins > 150 kDa. Membranes were blocked using 5% non-fat dry-milk in 0.1% TBST (0,05 mol/L Tris-HCL, 0,3 mol/L NaCl, 0,1% Tween 20) for 45 min at RT, then incubated ON at 4°C with primary antibodies as follow: anti-β-catenin diluted 1/1000 (Cell signaling, #2698), anti-tubulin diluted 1/500 (Sigma #I2G10), anti-CASK at 1 μg/ml final (K56A/50R; Addgene, #183002), anti-ARHGAP21 diluted 1/250 (Fisher scientific #16851715), anti-γ-catenin at 2 μg/ml final (clone 15F11; Antibodies #A249097), and anti-DLG1 at 8μg/ml final (Invitrogen #PA1-044). All primary antibodies dilutions were prepared in 5% non-fat dry milk in 0.1% TBST. HRP-conjugated secondary antibodies were diluted in 5% non-fat dry-milk in 0.1% TBST and used at 1/5000 for 2h at RT (anti-mouse Abcam #ab6728; anti-rabbit Abcam #ab6721). Blots were revealed using ECL (Roche #12015200001) or Femto ECL (GeneTex #GTX14698).

### SiRNA treatments

10^5^ cells were seeded on 22×22 mm glass coverslips. The day after, 30 pmol of double-stranded siRNAs were mixed separately with 4 µl of Jet Prime Reagent and 200 µl of Jet Prime Buffer. The mix was vortexed for 15 seconds and incubated at RT for 10 minutes before being added to the cells. After 24h, medium containing transfection reagents was washed with PBS and replaced with fresh medium. Cells were incubated for an additional 24h before use. SiRNA sequences are as follows:

FFL siRNA: 5’ rUrCrGrArArGrUrArCrUrCrArGrCrGrUrArArGTT 3’

APC siRNA: 5’ rGrCrArCrArArArGrCrUrGrUrUrGrArArUrUrUTT 3’

### Immunofluorescence

2.5×10^5^ cell were seeded on 22×22 mm glass coverslips. For HEK293 cells, coverslips were coated for 10 min with sterile 1% gelatin (Merck #VL655870) or with 0,01% poly-L-Lysine (Sigma Aldrich #P4707) before seeding cells. At 70% confluency, cells were fixed with 4% formaldehyde in PBS for 20 min at RT. Cells were washed twice with PBS then permeabilized with 0.1% triton (Thermo-scientific #28908) in PBS for 15 min at RT. Coverslips were washed twice with PBS and incubated with primary antibodies diluted in 0.1% BSA in PBS for 1h at RT. Primary antibodies were diluted as follow: APC 1/10 (Santa Cruz #sc-9998); β-catenin 1/400 (Cell signaling, #2698), JUP/γ-catenin 2 μg/ml (clone 15F11; Antibodies, A249097) and anti-ARHGAP21 diluted 1/250 (Fisher scientific #16851715). Cells were washed 3 times 10 min with PBS and then incubated for 1h at RT with fluorescent secondary antibodies from Jackson Immunoresearch, as follow: anti-mouse Cy3 diluted 1/1000 (#115-165-003); anti-rabbit Cy3 diluted 1/1000 (#711-166-152); anti-mouse FITC diluted 1/200 (#115-546-006); anti-rabbit FITC diluted 1/200 (#111-096-047) and anti-mouse Cy5 diluted 1/200 (#115-176-003). Cells were washed 3 times 10 min in PBS and mounted with Vectashield containing DAPI (Vector Laboratories).

### Single molecule fluorescent in situ hybridization (smFISH)

SmiFISH was performed as described previously (Tsanov et al., 2016). Briefly, 2.5×10^5^ cell were seeded on 22×22 mm glass coverslips. At 70% confluency, cells were fixed with 4% PFA in PBS for 20 min and permeabilized overnight with 70% ethanol at 4°C. Permeabilized cells were washed twice with PBS and incubated for 25 min at RT in a solution containing 15% formamide in 1xSSC. Unlabelled primary oligonucleotides were pre-hybridized with Cy3 or Cy5 labeled secondary oligonucleotides in a solution containing 100 mM NaCl, 50 mM tris-HCl, 10 mM MgCl_2_ (pH=7.9), for 3 min at 65°C to form a labeled duplex. Primary/secondary oligonucleotides ratios and sequences used for different mRNAs are provided in Table S3. Cells were hybridized overnight at 37°C with the duplex in the following mix: 15% Formamide, 0.34 μg/μl tRNA, 0.2 mg/ml RNase-free BSA, 2 mM VRC, 1x SSC and 10% dextran sulphate (MW 400-600 KDa). Cells were washed 3 times for 45 minutes in 15% formamide 1xSSC. Cells were then washed 3 times 1 minute with PBS. Finally, coverslips were mounted in Vectashield containing DAPI. With Cy5 labelled probes, the same protocol was used except that 0.1% Triton was added in the hybridization and washing buffers. SmFISH using directly conjugated Cy3 oligonucleotides was performed with probes hybridizing to GFP and IRES-NEO transcripts as described in (Chouaib et al., 2020).

### Fixed cell imaging

Images were acquired using an upright fluorescent wide-field microscope (Zeiss Axioimager Z3 apotome). Images were taken using 63X objective (Apochromat 1.46 NA oil), using Zen software. Multidimensional acquisition was used to acquire 3D Z-stacks in 4 channels: DAPI, GFP, Cy3 and Cy5. The distance between each image in the stack was set to 0.3 μm. Maximum projection was performed to transform Z-stacks into 2D images, using ImageJ, and Figures were mounted with Adobe Photoshop and Adobe Illustrator.

### Immunoprecipitation assays

3×10^6^ cell were seeded in 15 cm plate (5 plates/condition). At 70% confluency, cells were put on ice and gently washed twice with ice-cold PBS, then lysed with 1 ml HNTG lysis buffer (20 mM Hepes (pH=7,9), 150 mM NaCl, 1% Triton 100-X, 10% glycerol, 1 mM MgCl_2_, 1 mM EGTA) supplemented with 1x freshly prepared protease inhibitor cocktail (Roche #5056489001). Cells were incubated for 30 min on a tube roller at 4°C before being centrifuged for 10 min at 13,200 rpm at 4°C. Supernatants (1 ml) were incubated for 3h at 4°C with 50 μl M2 anti-Flag (Sigma-A2220) or GFP-TRAP agarose beads (Chromotek-gta-20). Wild-type untagged cells were used as a control. Beads were then washed 1 time with ice-cold HNTG and 3 times with ice-cold PBS. Immunoprecipitated proteins were eluted from the beads in 30 μl 1% SDS for 10 min at RT. After centrifugation, proteins were collected in a new tube and mixed with Laemmli (1X final).

### Label-free mass spectrometry

#### Sample preparation

Protein digestion was performed on S-TrapTM micro columns (Protifi) following the manufacturer’s instructions. In brief, protein extracts were diluted in 40 µL of final 5% SDS / 50 mM triethylammonium bicarbonate (TEAB), reduced with 20 mM dithiothreitol (DTT) and held for 10 min at 95°C. Samples were cooled to room temperature and alkylated with 40 mM iodoacetamide (IAA) for 30 min at room temperature in the dark. Samples were acidified with phosphoric acid at final concentration of 1.2% and diluted 6 times in S-Trap binding buffer (90% methanol / 100 mM TEAB). The resulting protein suspension was transferred to the S-Trap filter via centrifugation at 4000g for 1 min. Trapped proteins were washed three times with 150 µL S-Trap binding buffer. 1 µg of trypsin (Trypsin Glod, Promega) in 50 mM TEAB were added to the filter surface and incubated for 2 hours at 47°C. Tryptic peptides were eluted sequentially with 40 µl of 50 mM TEAB, 50 mM TEAB / 0.2% aqueous formic acid, and then with 50% of acetonitrinile (ACN) via centrifugation at 4000g. Eluted peptides were vacuum-dried.

#### Mass spectrometry

Experiments were performed on a Q Exactive HF mass spectrometer (Thermo Fisher Scientific) coupled with an Ultimate 3000 nano-HPLC (Thermo Fisher Scientific). Resulting peptide samples were solubilized in 0.05% trifluoroacetic acid (TFA) / 2% ACN, and were injected for desalting and pre-concentration on a PepMap®100 C18 precolumn (0.3 mm x 5 mm; ThermoFisher Scientific). Peptides separation was done on a 50 cm analytical reversed-phase column (75 mm inner diameter, Acclaim Pepmap 100® C18, Thermo Fisher Scientific) using a 123 min gradient of 2 to 40% of buffer B (80% ACN, 0.1% formic acid) and a flow rate of 300 nL/min.

MS/MS analyses were performed in a data-dependent mode. Full scans (375-1500 m/z) were acquired on the Orbitrap mass analyzer with a 60 000 resolution at 200 m/z. For the full scans, 3 e^6^ ions were accumulated within a maximum injection time of 60 ms and detected in the Orbitrap analyzer. The twelve most intense ions with charge states ≥ 2 were sequentially isolated to a target value of 1e5 with a maximum injection time of 100 ms and fragmented by HCD (Higher-energy collisional dissociation) in the collision cell (normalized collision energy of 28%) and detected in the Orbitrap analyzer at 30 000 resolution. First mass was set to 100 m/z.

#### Data analysis

Raw spectra were processed using the MaxQuant environment (Cox and Mann, 2008, v.2.0.3.0) and Andromeda for database search with label-free quantification (LFQ), match between runs and the iBAQ algorithm enabled (Cox et al., 2011). The MS/MS spectra were matched against the UniProt Reference proteome (Proteome ID UP000005640) of Homo sapiens (release 2023_03; https://www.uniprot.org/) and a homemade contaminant database. Enzyme specificity was set to trypsin/P, and the search included cysteine carbamidomethylation as a fixed modification and oxidation of methionine, and acetylation (protein N-term) as variable modifications. Up to two missed cleavages were allowed for protease digestion. FDR was set at 0.01 for peptides and proteins and the minimal peptide length at 7. A representative protein ID in each protein group was automatically selected using an in-house developed bioinformatics tool (Leading tool v3.5) as the best annotated protein in UniProtKB (reviewed entries rather than automatic ones; (Raynaud et al., 2018). Analysis of quantification data, statistical analyses and graphical representations were performed using Perseus (v1.6.15) (Rubinfeld et al., 1993). The mass spectrometry data have been deposited to the ProteomeXchange Consortium via the PRIDE partner repository with the dataset identifier PXD047379 (Fig. 4C and S3C), PXD047415 (Fig. 7D), PXD050354 (Fig. 4A), PXD047377 (Fig. 2D), PXD047369 (Fig. S3A) and PXD050179 (Fig. 2A).

### RNA immunoprecipitation

Immunoprecipitation was performed as above except that 3×10^6^ cells were used per condition (a 15 cm plate). Empty agarose beads (Chromotek-bab-20) were used as control. After 3h of immunoprecipitation and the 4 washes performed as above, RNeasy mini kit (Qiagen-74106) was used to purify immunoprecipitated RNAs. Beads were mixed with 200 μl of RLT lysis buffer, followed by 30 seconds vortex and 4 min incubation at RT. Beads and proteins were removed by centrifugation at 10,000g for 30 sec and the supernatant was transferred to a new 1.5 ml tube. One volume of 70% ethanol was added and mixed well by pipetting. The mixture was then transferred to a RNeasy column and centrifuged at 10,000g for 30 sec. RNAs were washed once with 700 μl of RW1 and 3 times with 700 μl of RPE buffers. Each wash step was followed by a quick spin at 10,000g for 30 sec except the last wash with RPE buffer which was centrifuged for 2 min. Columns were spined for 1 minute at 10,000g to remove excess liquid. Finally, the column was transferred to a new tube and RNAs were eluted in 30 μl RNase-free water. DNA contamination was eliminated by a routine DNase treatment using TURBO-DNA Free kit (Invitrogen-AM1907) as followed: 2 units of TURBO DNAse enzyme and 0.1 volume 10X TURBO DNAse Buffer were added to the sample in a final volume of 34 μl. The mix was incubated for 30 min at 37°C. After, 0.1 volume DNAse Inactivation Reagent was added to the mix and incubated for 5 min at 24°C. The tube was gently mixed 2-3 times during the incubation period. Samples were then centrifuged at 10,000xg for 2 minutes. The supernatant containing the RNAs was transferred to a new tube and samples were stored at -80°C.

### Sequencing and data analysis

SMART-Seq Stranded Kit (Takara Bio) was used to generate cDNA libraries for high-throughtput sequencing using NovaSeq 6000 Illumina. RNA and DNA libraries quality was subjected to quality control checks using a fragment bioanalyzer (Kit high sensivity NGS) and qPCR (ROCHE Light Cycler 480). Ribosomal cDNAs were cleaved using scZapR enzyme in the presence of scR-Probes designed against mammalian ribosomal RNAs. Libraries were then sequenced with NovaSeq Reagent Kit on a flow cell SP (single read -100 nucleotides) using the two-channel sequence by synthesis method. Before demultiplexing, sequences were mixed with the non-indexed PhiX sequences as an Illumina control. Demultiplexing was realized using Illumina bcl2fastq software (v2.20.0.422), thus effectively removing PhiX sequences. FastaQC software (v0.11.9) provided a tool to check reads quality after high-throughput sequencing and MultiQC program (V1.12) served to group data that emerges from all the samples. FastQ screen software served to detect contaminating genomes. Sequences were aligned on 15 species genomes, mycoplasma, and ribosomal RNAs using Bowtie2 software. No contaminations were detected.

TopHat2 software (v2.1.1) (APC and Axin experiments) or the HiSat2 (v2.2.1) (E-cadherin experiment) was used to align the obtained reads on the human genome (hg38 version) usinf a set of gene model annotations (gff file downloaded from UCSC on 2022-01-04). FeatureCounts (v2.0.3) software was utilized after to count reads per gene. All reads that overlap on two genes or show alignment at different positions on the genome were discarded. 80% of the counts showed unique alignments on the genome. Counts were normalized in each sample to eliminate differences due to manipulation. This was done by a normalization method known as relative log expression (RLE). Differential expression genes were identified using R (4.1.1) with packages EdgeR (v3.34.1) and DESeq2 (v1.32.0). The p-Value was considered significant if lower than 0.01. To calculate the reads per kilobase per million mapped reads (RPKM), the formula below was used:

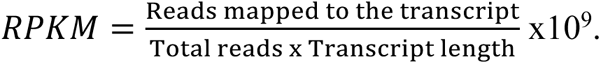

To calculate the transcripts per kilobase per million (TPM), the formula below was used:

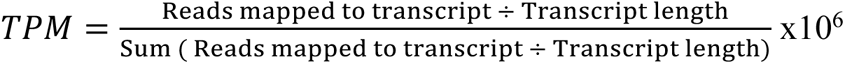

Sequencing data are accessible at GEO with the identifier GSE253965.

### Tissue smFISH

Mice experiments were performed according to the guidelines of the European Community (86/609/EEC) and the French National Committee (87/848) for care and use of laboratory animals, comply the ARRIVE guidelines and were approved by the French Ministry of Higher Education, Research and Innovation (reference APAFIS#18685) to be performed in the institute animal facility (agreement #F3417216). Mice were anesthetized with isoflurane then sacrificed via cervical dislocation. Colons from the caecum to the rectum were taken and flushed gently with PBS. Colons were then fixed for 3h at RT in 4% PFA in PBS before being transferred to 4% PFA 30% sucrose in PBS solution for an ON fixation at 4°C. The following day, the fixed tissue was embedded in a tissue embedding medium OCT (Fisher Scientific, # 23-730-571) and frozen by immersing it for 2-3 min in isopentane cooled to -80°C in liquid nitrogen. Sections of 6 mm thickness were generated using a Cryostat and collected on SuperFrost slides. Tissue sections were fixed again with 4%PFA in PBS solution for 20 min at RT then washed twice with 1xPBS. Sections were permeabilized with 70% ethanol ON at 4°C. For smFISH, the protocol was the same as for fixed cells. Only fresh blocs were used (<1 month old).

## Supplemental figures

**Figure S1: co-translational partners of APC (related to Figure 1)**

**A-**Schematic representation of the CRISPR repair cassette designed to insert 3xFLAG and mScarlet tags at the N-terminus of the APC protein. Puromycin: puromycin resistance gene; P2A: P2A self-cleaving peptides; HOM: homology arms, 3xFLAG: 3 repetitions of the FLAG tag.

**B-**Image showing an agarose gel loaded with the PCR products performed on both sides of the repair cassette integration site and obtained after amplification of genomic DNA. The edited allele of the clone N112 (APC KI N112) shows the expected amplicons of 975 bp and 788 bp on the 3’side and 5’side of the cassette, respectively. DNA ladder is loaded on the left showing the size of DNA markers.

**C-**Histogram showing transcript per kilobase million (TPKM) values of the most significant immunoprecipitated RNAs in 3xFLAG-mScarlet-APC HEK293 cells (-log_10_ (p-Value) ≥ 35.9). Experiments were done in duplicates. Data show means and standard deviation. Green: FLAG IP untreated; red: control IP untreated; blue: FLAG IP treated with puromycin; yellow: control IP treated with puromycin. Data show means and standard deviation.

**Figure S2: Axin1 co-translational partner (related to Figure 3)**

Histogram showing reads per kilobase million (RPKM) values of the most significant immunoprecipitated RNAs in 3xFLAG-eGFP-Axin1 HeLa H9 cells (-log_10_ (p-Value) ≥ 10.5). Experiments were done in duplicates. Green: FLAG IP untreated; red: control IP untreated; blue: FLAG IP treated with puromycin; yellow: control IP treated with puromycin. Data show means and standard deviation.

**Figure S3: Axin1 protein interactome with and without co-translational interactions (related to Figure 4)**

**A-**Volcano plot comparing label-free mass spectrometry analysis of 3xFLAG-eGFP-Axin1 IPs in HeLa cellsin puromycin treated *versus* untreated cells (30 min treatment at 100 μg/ml). The x axis displays the enrichment of proteins (log2 [ratio]) immunoprecipitated with anti-FLAG antibody conjugated beads (Flag IP) of 3xFLAG-eGFP-Axin1 cells treated with puromycin *versus* untreated cells. The y axis represents significance, displayed as -log10 (p-Value). Experiments were done in triplicates and statistical significance is considered for p-Value <0.01. Cells were treated with LG007 for 2h at 0.5 nM to stabilize Axin1 and obtain more materials in the pull-downs. The colored points in the plots represent different categories of proteins encoded by the identified mRNAs.

**B-**Western blot analysis showing β-catenin levels in H9 Flip-in cells treated with puromycin (100 μg/ml). Tubulin served as a loading control. Immunoblotted proteins are displayed on the left. Molecular protein weight markers size is indicated on the right. Lanes 2-3-4 were treated with LG007 for 2h at 0.5 nM.

**C**-Volcano plot representing label-free mass spectrometry analysis of eGFP-Axin1 immuno-precipitates in HEK293 cells. The x axis displays the enrichment of proteins (log2 [ratio]) immunoprecipitated with anti-GFP antibody conjugated beads (GFP-IP) in eGFP-Axin1 cells *versus* HEK293 wild-type cells used as control, both in normal conditions. The y axis represents significance, displayed as -log10 (p-Value). Experiments were done in triplicates and statistical significance is considered for p-Value <0.01.

**Figure S4: Composition of β-catenin mRNA foci in HeLa cells (related to Figure 6)**

**A-**Images are micrographs of HeLa β-catenin-GFP BAC cells. Left and red: Cy3 fluorescent signals corresponding to MYO5B, DLG1, AMER1 or ARHGAP21 mRNAs detected with smiFISH; middle and green: FITC signals corresponding to APC detected by immunofluorescence. For each cell, a zoom of the boxed area is shown below. Red arrows show APC foci and orange arrows show single mRNAs detected with MYO5B, DLG1, AMER1 or ARHGAP21 probes. DNA is stained with DAPI. Scale bar: 5 microns.

**B-**Images are micrographs of HeLa β-catenin GFP BAC cells. Left and red: Cy3 signals corresponding to MYO5B, DLG1, AMER1 or ARHGAP21 mRNAs detected with smiFISH; middle and green: Cy5 fluorescent signals corresponding to β-catenin mRNAs detected with smFISH. For each cell, a zoom of the boxed area is shown below. Red arrows show β-catenin mRNA foci and orange arrows show single mRNAs detected with MYO5B, DLG1, AMER1 or ARHGAP21 probes. DNA is stained with DAPI. Scale bar: 5 microns.

**Figure S5: β-catenin mRNA foci in wild-type mice colonic tissue (related to** Figure 6) Images are micrographs of mice wild-type colonic tissue with a crypt transversal section (upper) and longitudinal section (lower). Left and yellow represent Cy3 fluorescent signals corresponding to β-catenin mRNA detected with smiFISH, middle: autofluorescence signals captured with GFP channel. Red arrows show β-catenin mRNA foci and blue arrows show single molecules of β-catenin mRNA. DNA is stained with DAPI. Scale bar: 10 microns.

**Figure S6: Characterization of the HEK293 E-cadherin-GFP cell line (related to Figure 7)**

**A**-Images are micrographs of HEK293 cells expressing E-cadherin-GFP. Left and red: autofluorescence signals captured with the dsRed channel, middle and green: E-cadherin-eGFP signals. DNA is stained with DAPI. Scale bar: 10 microns.

**B-**Images are micrographs of HEK293 cells expressing E-cadherin-GFP and stained for β-catenin mRNA. Left and green: E-cadherin-eGFP signals, middle and red: Cy3 fluorescent signals corresponding to β-catenin mRNAs detected with smiFISH. For each cell, a zoom of the boxed area is shown below. Arrowheads show single molecules of β-catenin mRNA co-localizing with E-cadherin-eGFP. DNA is stained with DAPI. Scale bar: 5 microns.

## References

Aberle, H., Bauer, A., Stappert, J., Kispert, A., & Kemler, R. (1997). β-catenin is a target for the ubiquitin-proteasome pathway. EMBO Journal, 16(13), 3797–3804.

Albihlal, W. S., & Gerber, A. P. (2018). Unconventional RNA-binding proteins: an uncharted zone in RNA biology. In FEBS Letters (Vol. 592, Issue 17, pp. 2917–2931). Wiley Blackwell.

Behrens, J., von Kries, J. P., Kühl, M., Bruhn, L., Wedlich, D., Grosschedl, R., & Birchmeier, W. (1996). Functional interaction of beta-catenin with the transcription factor LEF-1. Nature, 382(6592), 638–642.

Bian, J., Dannappel, M., Wan, C., & Firestein, R. (2020a). Transcriptional Regulation of Wnt/β-Catenin Pathway in Colorectal Cancer. In Cells (Vol. 9, Issue 9). NLM (Medline).

Bian, J., Dannappel, M., Wan, C., & Firestein, R. (2020b). Transcriptional Regulation of Wnt/β-Catenin Pathway in Colorectal Cancer. In Cells (Vol. 9, Issue 9). NLM (Medline).

Buljan, M., Ciuffa, R., van Drogen, A., Vichalkovski, A., Mehnert, M., Rosenberger, G., Lee, S., Varjosalo, M., Pernas, L. E., Spegg, V., Snijder, B., Aebersold, R., & Gstaiger, M. (2020). Kinase Interaction Network Expands Functional and Disease Roles of Human Kinases. Molecular Cell, 79(3), 504–520.e9.

Chen, K. H., Boettiger, A. N., Moffitt, J. R., Wang, S., & Zhuang, X. (2015). Spatially resolved, highly multiplexed RNA profiling in sngle cells. Science, 348(6233).

Choi, H. J., Huber, A. H., & Weis, W. I. (2006). Thermodynamics of β-catenin-ligand interactions: The roles of the N- and C-terminal tails in modulating binding affinity. Journal of Biological Chemistry, 281(2), 1027–1038.

Chouaib, R., Safieddine, A., Pichon, X., Imbert, A., Kwon, O. S., Samacoits, A., Traboulsi, A. M., Robert, M. C., Tsanov, N., Coleno, E., Poser, I., Zimmer, C., Hyman, A., Le Hir, H., Zibara, K., Peter, M., Mueller, F., Walter, T., & Bertrand, E. (2020). A Dual Protein-mRNA Localization Screen Reveals Compartmentalized Translation and Widespread Co-translational RNA Targeting. Developmental Cell, 54(6), 773–791.e5.

Cselenyi, C. S., Jernigan, K. K., Tahinci, E., Thorne, C. A., Lee, L. A., & Lee, E. (2008). LRP6 transduces a canonical Wnt signal independently of Axin degradation by inhibiting GSK3’s phosphorylation of-catenin. Proceedings of the National Academy of Sciences of the United States of America vol. 105,23 (2008): 8032–7.

Cox, J., & Mann, M. (2008). MaxQuant enables high peptide identification rates, individualized p.p.b.-range mass accuracies and proteome-wide protein quantification. Nature biotechnology, 26(12), 1367– 1372.

Cox, J., Neuhauser, N., Michalski, A., Scheltema, R. A., Olsen, J. V., & Mann, M. (2011). Andromeda: a peptide search engine integrated into the MaxQuant environment. Journal of proteome research, 10(4), 1794–1805.

Drees, F., Pokutta, S., Yamada, S., Nelson, W. J., & Weis, W. I. (2005). α-catenin is a molecular switch that binds E-cadherin-β-catenin and regulates actin-filament assembly. Cell, 123(5), 903–915.

Faux, M. C., Coates, J. L., Catimel, B., Cody, S., Clayton, A. H. A., Layton, M. J., & Burgess, A. W. (2008). Recruitment of adenomatous polyposis coli and β-catenin to axin-puncta. Oncogene, 27(44), 5808–5820.

Gerashchenko, M. V., Peterfi, Z., Yim, S. H., & Gladyshev, V. N. (2021). Translation elongation rate varies among organs and decreases with age. Nucleic Acids Research, 49(2), E9.

Gergen, J. P., & Wieschaus, E. F. (1986). Localized requirements for gene activity in segmentation of Drosophila embryos: analysis of armadillo, fused, giant and unpaired mutations in mosaic embryos. In Arch Dev Biol (Vol. 195).

Goto, T., Matsuzawa, J., Iemura, S. I., Natsume, T., & Shibuya, H. (2016). WDR26 is a new partner of Axin1 in the canonical Wnt signaling pathway. FEBS Letters, 590(9), 1291–1303.

Graham, T. A., Clements, W. K., Kimelman, D., & Xu, W. (2002). The crystal structure of the beta-catenin/ICAT complex reveals the inhibitory mechanism of ICAT. Molecular cell, 10(3), 563–571.

Grainger, S., & Willert, K. (2018). Mechanisms of Wnt signaling and control. In Wiley Interdisciplinary Reviews: Systems Biology and Medicine (Vol. 10, Issue 5).

Groden, J., Thliveris, ’t Wade Samowitz, A., Carlson, M., Gelbert’t, L., Albertsen, H., Joslyn, G., Stevens, J., Spirio, L., Robertson’t, M., Sargeant, L., Krapcho,’, K., Wolff, E., Burt, R., Hughes, J. P., Warrington, J., Mcpherson, J., Wasmuth, J., Le Paslier, D., Abderrahim, H., … White’?, R. (1991). Identification and Characterization of the Familial Adenomatous Polyposis Coli Gene. In Cell (Vol. 66).

Grohmann, A., Tanneberger, K., Alzner, A., Schneikert, J., & Behrens, J. (2007). AMER1 regulates the distribution of the tumor suppressor APC between microtubules and the plasma membrane. Journal of Cell Science, 120(21), 3738–3747.

Grossmann, T. N., Yeh, J. T. H., Bowman, B. R., Chu, Q., Moellering, R. E., & Verdine, G. L. (2012). Inhibition of oncogenic Wnt signaling through direct targeting of β-catenin. Proceedings of the National Academy of Sciences of the United States of America, 109(44), 17942–17947.

Ha, et al. (2004). Mechanism of Phosphorylation-Dependent Binding of APC to-Catenin and Its Role in-Catenin Degradation. Molecular cell vol. 15,4 (2004): 511–21.

Hart, M. J., De Los Santos, R., Albert, I. N., Rubinfeld, B., & Polakis, P. (1998). Downregulation of b-catenin by human Axin and its association with the APC tumor suppressor, b-catenin and GSK3b. Current biology: CB vol. 8,10 (1998): 573–81.

Hein, M. Y., Hubner, N. C., Poser, I., Cox, J., Nagaraj, N., Toyoda, Y., Gak, I. A., Weisswange, I., Mansfeld, J., Buchholz, F., Hyman, A. A., & Mann, M. (2015). A Human Interactome in Three Quantitative Dimensions Organized by Stoichiometries and Abundances. Cell, 163(3), 712–723.

Hentze, M. W., Castello, A., Schwarzl, T., & Preiss, T. (2018). A brave new world of RNA-binding proteins. In Nature Reviews Molecular Cell Biology (Vol. 19, Issue 5, pp. 327–341). Nature Publishing Group.

Hiilsken, J., Birchmeier, W., & Behrens, J. (1994). E-Cadherin and APC Compete for the Interaction with-Catenin and the Cytoskeleton. The Journal of cell biology vol. 127,6 Pt 2 (1994): 2061–9.

Hirschberg, K., Miller, C. M., Ellenberg, J., Presley, J. F., Siggia, E. D., Phair, R. D., & Lippincott-Schwartz, J. (1998). Kinetic Analysis of Secretory Protein Traffic and Characterization of Golgi to Plasma Membrane Transport Intermediates in Living Cells. In The Journal of Cell Biology (Vol. 143, Issue 6).

Huber, A. H., & Weis, W. I. (2001). The structure of the beta-catenin/E-cadherin complex and the molecular basis of diverse ligand recognition by beta-catenin. Cell, 105(3), 391–402.

Ikeda, S., Kishida, S., Yamamoto, H., Murai, H., Koyama, S., & Kikuchi, A. (1998a). Axin, a negative regulator of the Wnt signaling pathway, forms a complex with GSK-3β and β-catenin and promotes GSK-3β-dependent phosphorylation of β-catenin. EMBO Journal, 17(5), 1371–1384.

Ikeda, S., Kishida, S., Yamamoto, H., Murai, H., Koyama, S., & Kikuchi, A. (1998b). Axin, a negative regulator of the Wnt signaling pathway, forms a complex with GSK-3β and β-catenin and promotes GSK-3β-dependent phosphorylation of β-catenin. In The EMBO Journal (Vol. 17, Issue 5).

Iwao, K., Nakamori, S., Kameyama, M., Imaoka, S., Kinoshita, M., Fukui, T., Ishiguro, S., Nakamura, Y., & Miyoshi, Y. (1998). Activation of the beta-catenin gene by interstitial deletions involving exon 3 in primary colorectal carcinomas without adenomatous polyposis coli mutations. Cancer research, 58(5), 1021–1026.

Jia, L., Miao, C., Cao, Y., & Duan, E. kui. (2008). Effects of Wnt proteins on cell proliferation and apoptosis in HEK293 cells. Cell Biology International, 32(7), 807–813.

Kinzler, K. W., Nilbert, M. C., Su, L.-K., Vogelstein, B., Bryan, T. M., Levy, D. B., Smith, K. J., Preisinger, A. C., Hedge, P., Mckechnie, D., Finniear, R., Markham, A., Groffen, J., Boguski, M. S., Altschul, S. F., Horii, A., Ando, H., Miyoshi, Y., Miki, Y., … Nakamura, Y. (n.d.). Identification of FAP Locus Genes from Chromosome 5q21. In Source: Science, New Series (Vol. 253, Issue 5020).

Kodama, S., Ikeda, S., Asahara, T., Kishida, M., & Kikuchi, A. (1999). Axin directly interacts with plakoglobin and regulates its stability. Journal of Biological Chemistry, 274(39), 27682–27688.

Kolligs, F. T., Kolligs, B., Hajra, K. M., Hu, G., Tani, M., Cho, K. R., & Fearon, E. R. (2000). Catenin is regulated by the APC tumor suppressor and its oncogenic activity is distinct from that of-catenin. Genes & development vol. 14,11 (2000): 1319-31.

Korinek, V., Barker, N., Moerer, P., van Donselaar, E., Huls, G., Peters, P. J., & Clevers, H. (1998). Depletion of epithelial stem-cell compartments in the small intestine of mice lacking Tcf-4. Nature genetics, 19(4), 379–383.

Kwong, L. N., & Dove, W. F. (2009). APC and its modifiers in colon cancer NIH Public Access. In Adv Exp Med Biol (Vol. 656).

Liang, X., Men, Q. L., Li, Y. W., Li, H. C., Chong, T., & Li, Z. L. (2017). Silencing of armadillo repeat-containing protein 8 (ARMc8) inhibits TGF-β-induced EMT in bladder carcinoma UMUC3 Cells. Oncology Research, 25(1), 99–105.

Liu, C., Kato, Y., Zhang, Z., Do, M., Yankner, B. A., & He, X. I. (1999). catenin phosphorylation-degradation and regulates Xenopus axis formation. In Developmental Biology-Trcp couples (Vol. 96).

Liu, C., Li, Y., Semenov, M., Han, C., Baeg, G.-H., Tan, Y., Zhang, Z., Lin, X., & He, X. (2002). Control of-Catenin Phosphorylation/Degradation by a Dual-Kinase Mechanismcatenin phosphorylation controls. In genes (Vol. 108).

Lock, J. G., Hammond, L. A., Houghton, F., Gleeson, P. A., & Stow, J. L. (2005). E-cadherin transport from the trans-Golgi network in tubulovesicular carriers is selectively regulated by Golgin-97. Traffic, 6(12), 1142–1156.

MacDonald, B. T., Tamai, K., & He, X. (2009). Wnt/β-Catenin Signaling: Components, Mechanisms, and Diseases. In Developmental Cell (Vol. 17, Issue 1, pp. 9–26).

Matsumine, A., Ogai, A., Senda, T., Okumura, N., Satoh, K., Baeg, G. H., Kawahara, T., Kobayashi, S., Okada, M., Toyoshima, K., & Akiyama, T. (1996). Binding of APC to the human homolog of the Drosophila discs large tumor suppressor protein. Science (New York, N.Y.), 272(5264), 1020–1023.

Mccrea, P. D., Turck, C. W., & Gumbiner, B. (1991). A Homolog of the armadillo Protein in Drosophila (Plakoglobin) Associated with E-Cadherin. Science (New York, N.Y.) vol. 254,5036 (1991): 1359-61.

McEwen, A. E., Maher, M. T., Mo, R., & Gottardi, C. J. (2014). E-cadherin phosphorylation occurs during its biosynthesis to promote its cell surface stability and adhesion. Molecular Biology of the Cell, 25(16), 2365–2374.

Mills, K. M., Brocardo, M. G., & Henderson, B. R. (2016). APC binds the miro/Milton motor complex to stimulate transport of mitochondria to the plasma membrane. Molecular Biology of the Cell, 27(3), 466–482.

Miyoshi, Y., Ando, H., Nagase, H., Nishisho, I., Horii, A., Miki, Y., Morit, T., Utsunomiyat, J., Baba, S., Petersen, G., Hamilton, S. R., Kinzler, K. W., Vogelstein, B., & Nakamura, Y. (1992). Germ-line mutations of the APC gene in 53 familial adenomatous polyposis patients (Vol. 89).

Miyoshi, Y., Iwao, K., Nagasawa, Y., Aihara, T., Sasaki, Y., Imaoka, S., Murata, M., Shimano, T., & Nakamura, Y. (1998). Activation of the beta-catenin gene in primary hepatocellular carcinomas by somatic alterations involving exon 3. Cancer research, 58(12), 2524–2527.

Morin, P. J., Sparks, A. B., Korinek, V., Barker, N., Clevers, H., Vogelstein, B., & Kinzler, K. W. (1997). Activation of beta-catenin-Tcf signaling in colon cancer by mutations in beta-catenin or APC. Science (New York, N.Y.), 275(5307), 1787–1790.

Nakanishi, H., Sawada, T., Kaizaki, Y., Ota, R., Suzuki, H., Yamamoto, E., Aoki, H., Eizuka, M., Hasatani, K., Takahashi, N., Inagaki, S., Ebi, M., Kato, H., Kubota, E., Kataoka, H., Takahashi, S., Tokino, T., Minamoto, T., Sugai, T., & Sasaki, Y. (2020). Significance of gene mutations in the Wnt signaling pathway in traditional serrated adenomas of the colon and rectum. PLoS ONE, 15(2).

Nong, J., Kang, K., Shi, Q., Zhu, X., Tao, Q., & Chen, Y. G. (2021). Phase separation of Axin organizes the β-catenin destruction complex. Journal of Cell Biology, 220(4).

Nusse, R., & Varmus, H. E. (1982). Many Tumors Induced by the Mouse Mammary Tumor Virus Contain a Provirus Integrated in the Same Region of the Host Genome. In Cell (Vol. 31).

Ozawa, M., Baribault, H., & Kemler, R. (1989). The cytoplasmic domain of the cell adhesion molecule uvomorulin associates with three independent proteins structurally related in different species. In The EMBO Journal (Vol. 8, Issue 6).

Peifer, M., & Yap, A. S. (2003). Traffic control: p120-catenin acts as a gatekeeper to control the fate of classical cadherins in mammalian cells. The Journal of cell biology, 163(3), 437–440.

Piao, S., Lee, S. H., Kim, H., Yum, S., Stamos, J. L., Xu, Y., Lee, S. J., Lee, J., Oh, S., Han, J. K., Park, B. J., Weis, W. I., & Ha, N. C. (2008). Direct inhibition of GSK3β by the phosphorylated cytoplasmic domain of LRP6 in Wnt/β-catenin signaling. PLoS ONE, 3(12).

Polakis P. (2007). The many ways of Wnt in cancer. Current opinion in genetics & development, 17(1), 45–51.

Preitner, N., Quan, J., Nowakowski, D. W., Hancock, M. L., Shi, J., Tcherkezian, J., Young-Pearse, T. L., & Flanagan, J. G. (2014). APC is an RNA-binding protein, and its interactome provides a link to neural development and microtubule assembly. Cell, 158(2), 368–382.

Pronobis, M. I., Rusan, N. M., & Peifer, M. (2015). A novel GSK3-regulated APC:Axin interaction regulates Wnt signaling by driving a catalytic cycle of efficient βcatenin destruction. eLife vol. 4 e08022.

Raynaud, F., Homburger, V., Séveno, M., Vigy, O., Moutin, E., Fagni, L., Perroy, J., & Raynaud, F. (2018). SNAP23-Kif5 complex controls mGlu1 receptor trafficking. Journal of Molecular Cell Biology, 10(5), 423–436.

Rubinfeld, B., Albert, I., Porfiri, E., Fiol, C., Munemitsu, S., & Polakis, P. (1996). Binding of GSK3β to the APC-β-Catenin Complex and Regulation of Complex Assembly. In Source: Science, New Series (Vol. 272, Issue 5264).

Rubinfeld, B., Souza, B., Albert, I., Munemitsu, S., & Polakisz, P. (1995). THE JOURNAL OF BIOLOGICAL CHEMISTRY The APC Protein and E-cadherin Form Similar but Independent Complexes with o-Catenin, Il-Catenin, and Plakoglobin* (Vol. 270, Issue 10).

Rubinfeld, B., Souza, B., Albert, I., Müller, O., Chamberlain, S. H., Masiarz, F. R., Munemitsu, S., & Polakis, P. (1993). Association of the APC gene product with beta-catenin. Science (New York, N.Y.), 262(5140), 1731–1734.

Sato, A., Shimizu, M., Goto, T., Masuno, H., Kagechika, H., Tanaka, N., & Shibuya, H. (2020). WNK regulates Wnt signalling and β-Catenin levels by interfering with the interaction between β-Catenin and GID. Communications Biology, 3(1).

Schaefer, K. N., & Peifer, M. (2019). Wnt/Beta-Catenin Signaling Regulation and a Role for Biomolecular Condensates. In Developmental Cell (Vol. 48, Issue 4, pp. 429–444). Cell Press.

Sollerbrant, K., Chinnadurai, G., & Svensson, C. (1996). The CtBP binding domain in the adenovirus E1A protein controls CR1-dependent transactivation. In Nucleic Acids Research (Vol. 24, Issue 13).

Sousa, S., Cabanes, D., Archambaud, C., Colland, F., Lemichez, E., Popoff, M., Boisson-Dupuis, S., Gouin, E., Lecuit, M., Legrain, P., & Cossart, P. (2005). ARHGAP10 is necessary for α-catenin recruitment at adherens junctions and for Listeria invasion. Nature Cell Biology, 7(10), 954–960.

Stamos, J. L., & Weis, W. I. (2013). The β-catenin destruction complex. In Cold Spring Harbor Perspectives in Biology (Vol. 5, Issue 1).

Trcek, T., Douglas, T. E., Grosch, M., Yin, Y., Eagle, W. V. I., Gavis, E. R., Shroff, H., Rothenberg, E., & Lehmann, R. (2020). Sequence-Independent Self-Assembly of Germ Granule mRNAs into Homotypic Clusters. Molecular Cell, 78(5), 941–950.e12.

Trendel, J., Schwarzl, T., Horos, R., Prakash, A., Bateman, A., Hentze, M. W., & Krijgsveld, J. (2019). The Human RNA-Binding Proteome and Its Dynamics during Translational Arrest. Cell, 176(1–2), 391–403.e19.

Tsanov, N., Samacoits, A., Chouaib, R., Traboulsi, A. M., Gostan, T., Weber, C., Zimmer, C., Zibara, K., Walter, T., Peter, M., Bertrand, E., & Mueller, F. (2016). SmiFISH and FISH-quant - A flexible single RNA detection approach with super-resolution capability. Nucleic Acids Research, 44(22).

Yamamoto, H., Kishida, S., Kishida, M., Ikeda, S., Takada, S., & Kikuchi, A. (1999). Phosphorylation of Axin, a Wnt Signal Negative Regulator, by Glycogen Synthase Kinase-3 Regulates Its Stability. The Journal of biological chemistry vol. 274,16 (1999): 10681-4.

