## Supplementary figures for "Co-translational sorting enables a single mRNA to generate distinct polysomes with different localizations and protein fates"

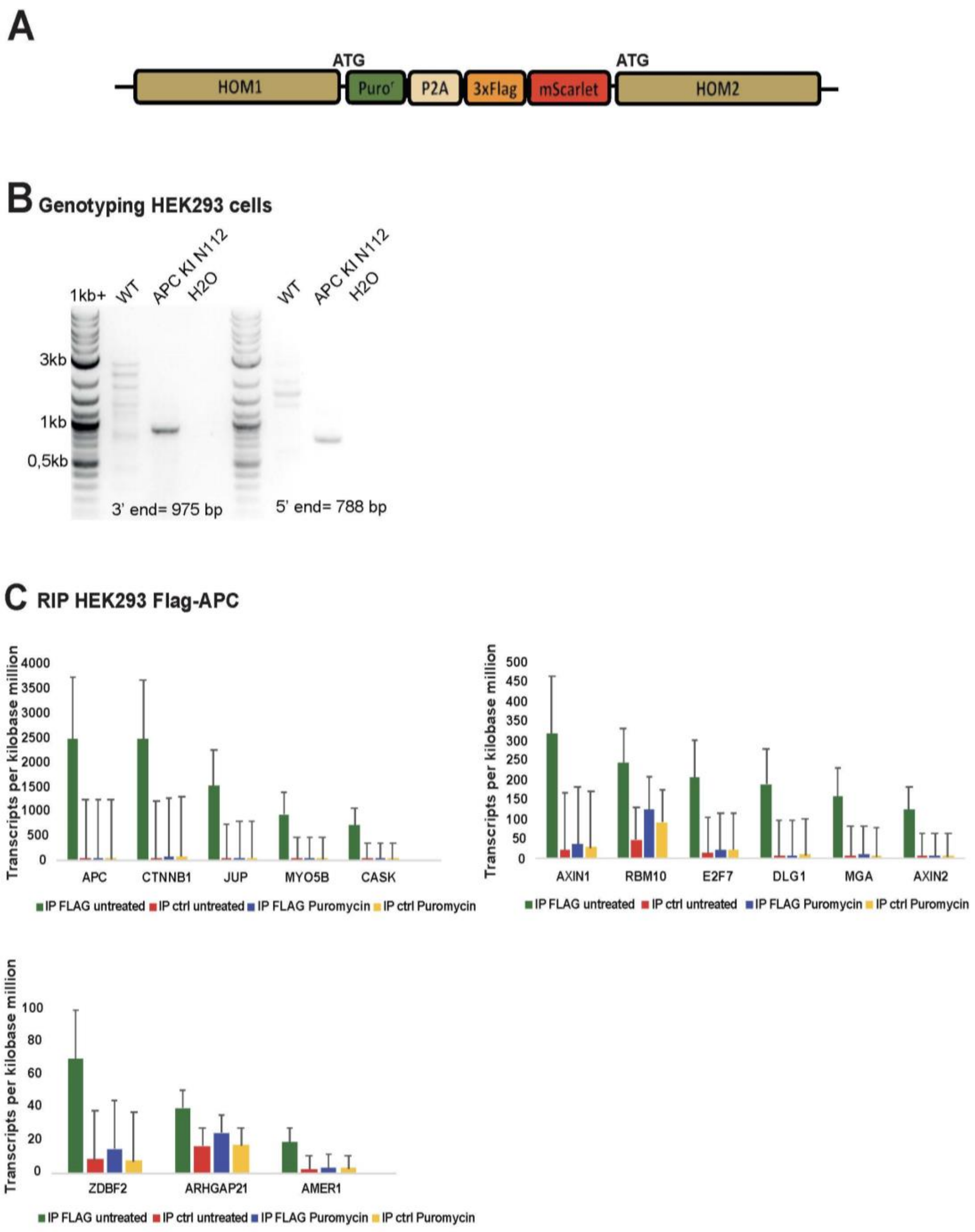


**Figure S1: co-translational partners of APC (related to Figure 1)**

**A-**Schematic representation of the CRISPR repair cassette designed to insert 3xFLAG and mScarlet tags at the N-terminus of the APC protein. Puromycin: puromycin resistance gene; P2A: P2A self-cleaving peptides; HOM: homology arms, 3xFLAG: 3 repetitions of the FLAG tag.

**B-**Image showing an agarose gel loaded with the PCR products performed on both sides of the repair cassette integration site and obtained after amplification of genomic DNA. The edited allele of the clone N112 (APC KI N112) shows the expected amplicons of 975 bp and 788 bp on the 3'side and 5'side of the cassette, respectively. DNA ladder is loaded on the left showing the size of DNA markers.

**C-**Histogram showing transcript per kilobase million (TPKM) values of the most significant immunoprecipitated RNAs in 3xFLAG-mScarlet-APC HEK293 cells ($-\log_{10}$ (p-Value) $\geq 35.9$). Experiments were done in duplicates. Data show means and standard deviation. Green: FLAG IP untreated; red: control IP untreated; blue: FLAG IP treated with puromycin; yellow: control IP treated with puromycin. Data show means and standard deviation.

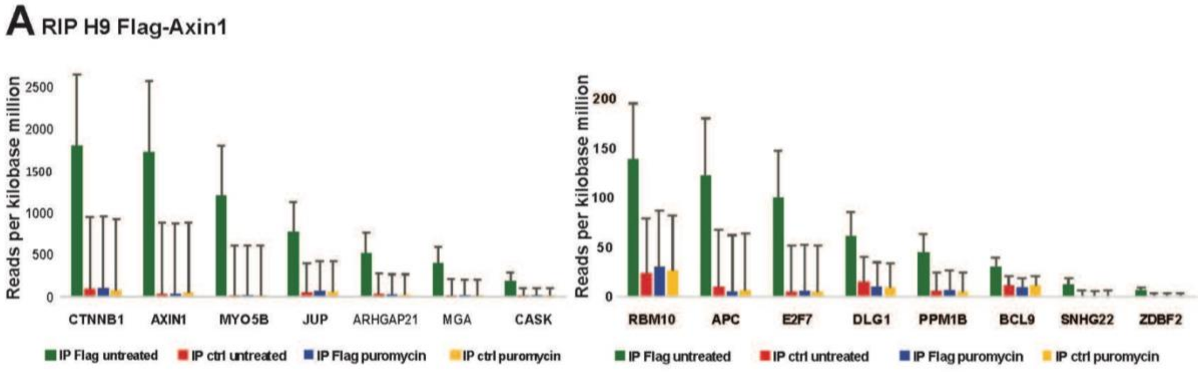


**Figure S2: Axin1 co-translational partner (related to Figure 3)**

Histogram showing reads per kilobase million (RPKM) values of the most significant immunoprecipitated RNAs in 3xFLAG-eGFP-Axin1 HeLa H9 cells (-$\log_{10}$ (p-Value) $\geq$ 10.5). Experiments were done in duplicates. Green: FLAG IP untreated; red: control IP untreated; blue: FLAG IP treated with puromycin; yellow: control IP treated with puromycin. Data show means and standard deviation.

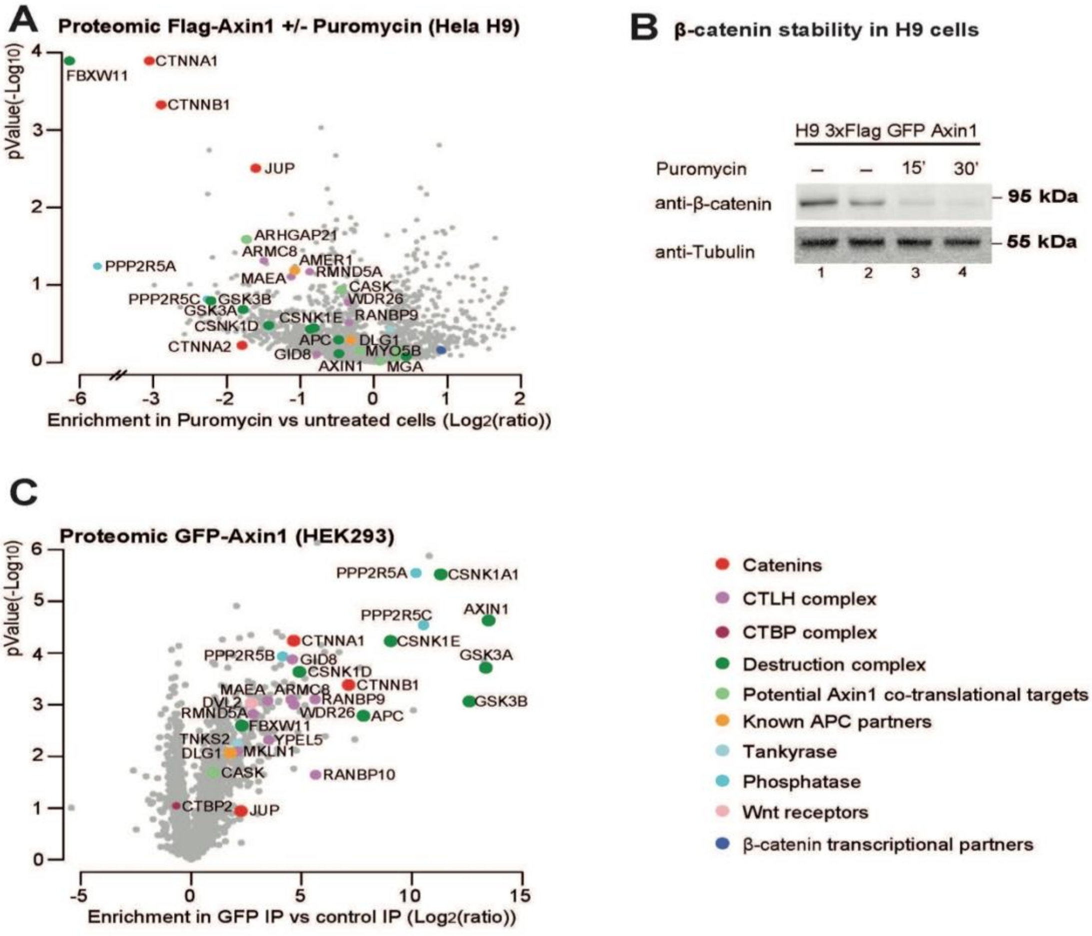


**Figure S3: Axin1 protein interactome with and without co-translational interactions (related to Figure 4)**

**A-**Volcano plot comparing label-free mass spectrometry analysis of 3xFLAG-eGFP-Axin1 IPs in HeLa cellsin puromycin treated *versus* untreated cells (30 min treatment at 100 µg/ml). The x axis displays the enrichment of proteins (log2 [ratio]) immunoprecipitated with anti-FLAG antibody conjugated beads (Flag IP) of 3xFLAG-eGFP-Axin1 cells treated with puromycin *versus* untreated cells. The y axis represents significance, displayed as -log10 (p-Value). Experiments were done in triplicates and statistical significance is considered for p-Value <0.01. Cells were treated with LG007 for 2h at 0.5 nM to stabilize Axin1 and obtain more materials in the pull-downs. The colored points in the plots represent different categories of proteins encoded by the identified mRNAs.

**B**-Western blot analysis showing β-catenin levels in H9 Flip-in cells treated with puromycin (100 μg/ml). Tubulin served as a loading control. Immunoblotted proteins are displayed on the left. Molecular protein weight markers size is indicated on the right. Lanes 2-3-4 were treated with LG007 for 2h at 0.5 nM.

**C**-Volcano plot representing label-free mass spectrometry analysis of eGFP-Axin1 immuno-precipitates in HEK293 cells. The x axis displays the enrichment of proteins (log2 [ratio]) immunoprecipitated with anti-GFP antibody conjugated beads (GFP-IP) in eGFP-Axin1 cells *versus* HEK293 wild-type cells used as control, both in normal conditions. The y axis represents significance, displayed as -log10 (p-Value). Experiments were done in triplicates and statistical significance is considered for p-Value <0.01.

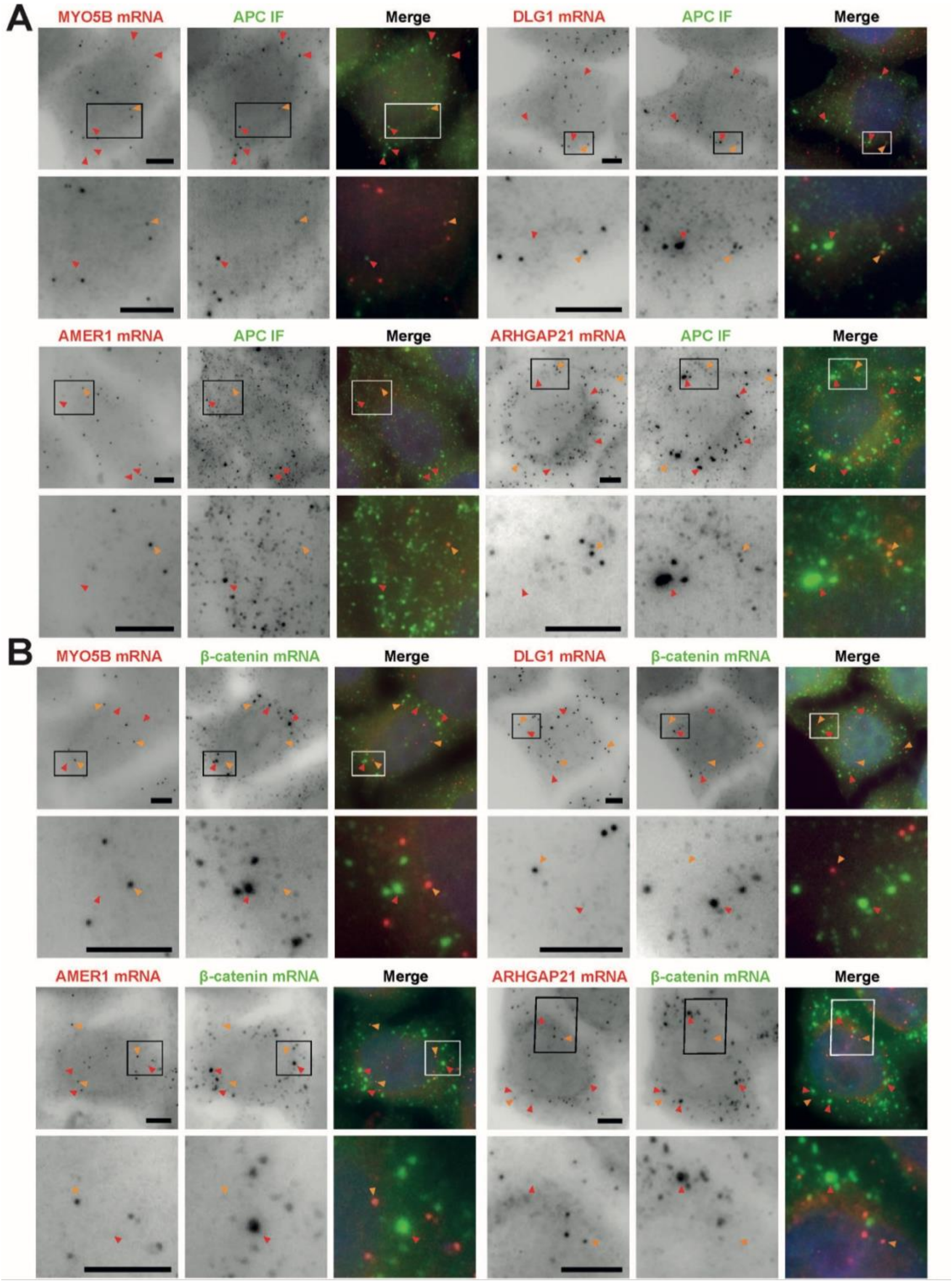


**Figure S4: Composition of β-catenin mRNA foci in HeLa cells (related to Figure 6)**

**A-**Images are micrographs of HeLa β-catenin-GFP BAC cells. Left and red: Cy3 fluorescent signals corresponding to MYO5B, DLG1, AMER1 or ARHGAP21 mRNAs detected with smiFISH; middle and green: FITC signals corresponding to APC detected by immunofluorescence. For each cell, a zoom of the boxed area is shown below. Red arrows show APC foci and orange arrows show single mRNAs detected with MYO5B, DLG1, AMER1 or ARHGAP21 probes. DNA is stained with DAPI. Scale bar: 5 microns.

**B-**Images are micrographs of HeLa β-catenin GFP BAC cells. Left and red: Cy3 signals corresponding to MYO5B, DLG1, AMER1 or ARHGAP21 mRNAs detected with smiFISH; middle and green: Cy5 fluorescent signals corresponding to β-catenin mRNAs detected with smFISH. For each cell, a zoom of the boxed area is shown below. Red arrows show β-catenin mRNA foci and orange arrows show single mRNAs detected with MYO5B, DLG1, AMER1 or ARHGAP21 probes. DNA is stained with DAPI. Scale bar: 5 microns.

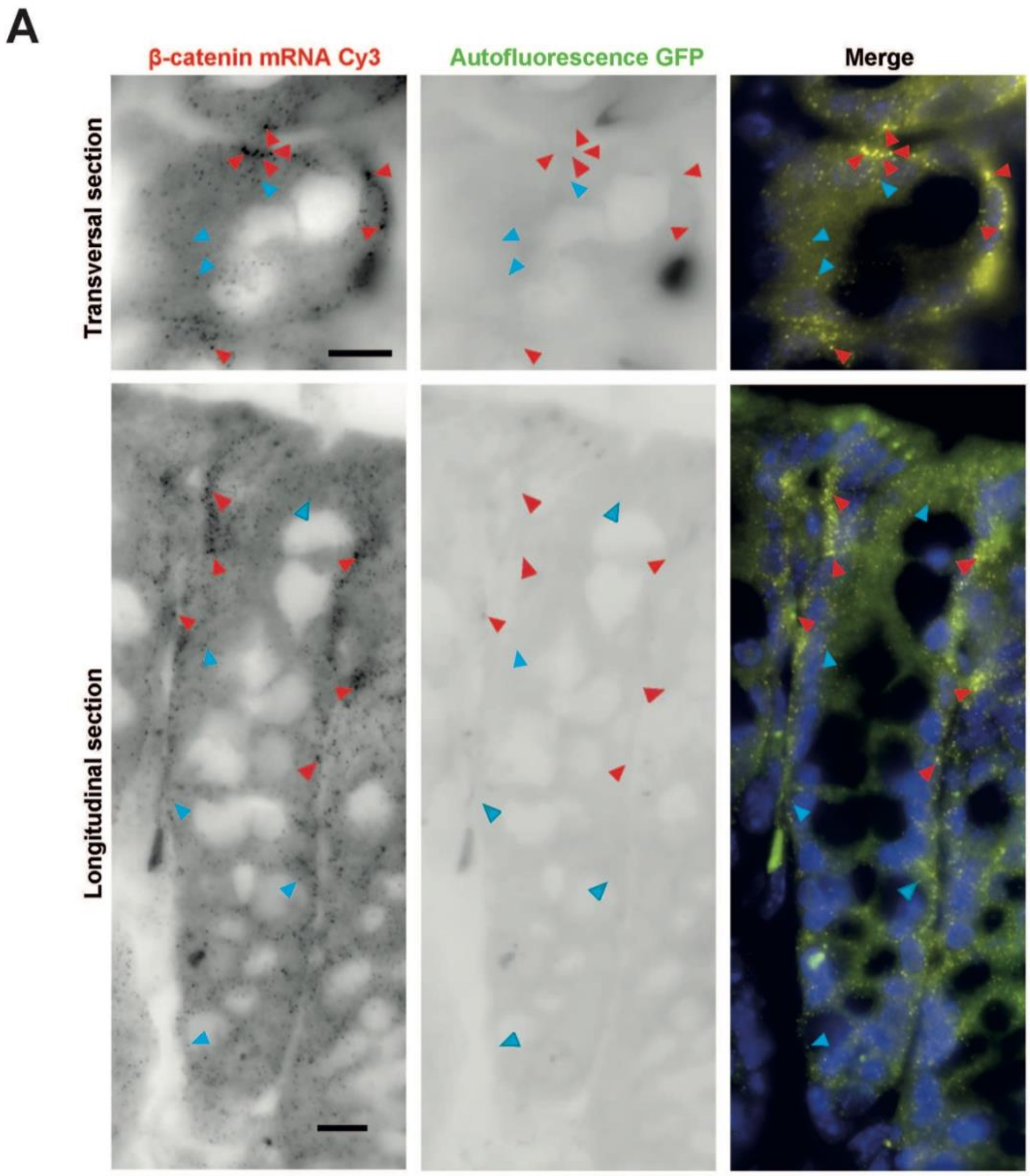


**Figure S5: β-catenin mRNA foci in wild-type mice colonic tissue (related to Figure 6)**

Images are micrographs of mice wild-type colonic tissue with a crypt transversal section (upper) and longitudinal section (lower). Left and yellow represent Cy3 fluorescent signals corresponding to β-catenin mRNA detected with smiFISH, middle: autofluorescence signals captured with GFP channel. Red arrows show β-catenin mRNA foci and blue arrows show single molecules of β-catenin mRNA. DNA is stained with DAPI. Scale bar: 10 microns.

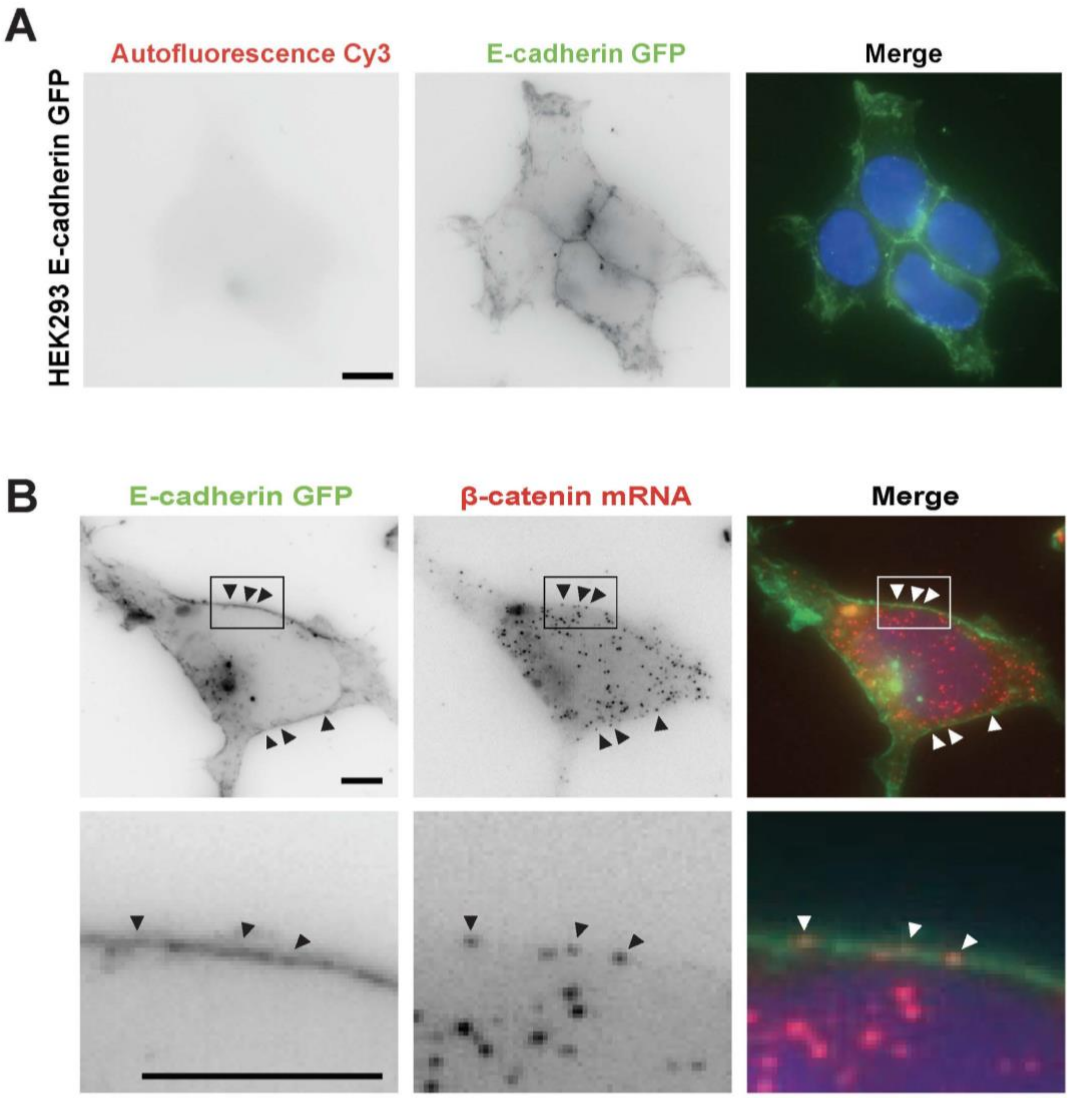


**Figure S6: Characterization of the HEK293 E-cadherin-GFP cell line (related to Figure 7)**

**A**-Images are micrographs of HEK293 cells expressing E-cadherin-GFP. Left and red: autofluorescence signals captured with the dsRed channel, middle and green: E-cadherin-eGFP signals. DNA is stained with DAPI. Scale bar: 10 microns.

**B**-Images are micrographs of HEK293 cells expressing E-cadherin-GFP and stained for β-catenin mRNA. Left and green: E-cadherin-eGFP signals, middle and red: Cy3 fluorescent signals corresponding to β-catenin mRNAs detected with smiFISH. For each cell, a zoom of the boxed area is shown below. Arrow heads show single molecules of β-catenin mRNA co-localizing with E-cadherin-eGFP. DNA is stained with DAPI. Scale bar: 5 microns.
